# When is microbial cross-feeding evolutionarily stable?

**DOI:** 10.1101/2025.05.16.654511

**Authors:** Jamie Alcira Lopez, Bo Liu, Zhiyuan Li, Mohamed S. Donia, Ned S. Wingreen

## Abstract

Cross-feeding, a phenomenon in which organisms share metabolites, is frequently observed in microbial communities across the natural world. One of the most common forms is waste-product cross-feeding, a unidirectional interaction in which the waste products of one microbe support the growth of another. Despite its ubiquity, it is not well-understood why waste-product cross-feeding persists when a single organism could in principle perform both the producer and consumer role. To address this question, we first analyze cross-feeding evolution in a minimal model of microbial metabolism. The model describes multi-step extraction of energy from a substrate in a simple but thermodynamically correct formulation. Surprisingly, we find that cross-feeding is never evolutionarily stable in this model. By analyzing models with more complex growth functions, we identify a novel mechanism for the evolutionary stability of waste-product cross-feeding, namely, generalized intracellular metabolite toxicity. Such toxicity arises because, in excess, the same intracellular metabolites that cells require for metabolism can be detrimental to growth (e.g., due to osmotic stress). We show that some but not all forms of such toxicity can lead to evolutionarily stable consortia of microbes that cross-feed waste products. This stability results from the potential of such consortia to divide the burden of toxic metabolites among a larger population, allowing them to perform their collective metabolism more efficiently than non-cross-feeders. More generally, we predict that growth penalties that scale nonlinearly with intracellular metabolite levels promote cross-feeding. We find that this mechanism for cross-feeding evolutionary stability implies nontrivial population dynamics, such as a discontinuity in population biomass at the onset of cross-feeding.

**Significance statement:** The chemical reactions performed by microbes have large impacts on our world: from nitrification within the nitrogen cycle to the breakdown of fiber in animal digestive tracts. A striking commonality in many of these processes is that the chemical reactions are a collective effort, with the complete reaction subdivided between many microbes. This phenomenon is known as cross-feeding, and its origins are poorly understood. Understanding the eco-evolutionary forces promoting cross-feeding have the potential to not only enhance our understanding of natural ecosystems, but also improve our ability to engineer such distributed reactions in biotechnology. Here, we develop mathematical theory for the evolution of a common type of cross-feeding and provide predictions for what metabolic and environmental conditions promote this behavior.

## Introduction

Microbial communities are found nearly everywhere on our planet, in environments ranging from the human gut (*1*) to deep-sea hydrothermal vents (*2*). One of the most striking properties of these communities is the ubiquity of chemical exchange: despite being nominally selfish organisms, microbes engage in prolific sharing of useful chemical compounds. This exchange is known as cross-feeding, and has been shown to influence a number of ecosystem properties. For example, cross-feeding in pathogen communities can enhance virulence (*3*), while in biotechnological settings disruption of cross-feeding can lead to failure of wastewater treatment (*4*). Recent experimental and theoretical ecology work suggests that cross-feeding may underpin the diversity and structure of microbial communities in a predictable manner (*5*–*7*). Thus, cross-feeding needs to be understood to predict and engineer microbial communities.

One of the most geochemically and biotechnologically important forms of cross-feeding is waste-product cross-feeding, in which some microbes derive energy from the waste products of other microbe’s metabolism. Examples include “anaerobic digestion”, in which many classes of organism cooperatively degrade complex organic polymers (*8, 9*) and nitrification, in which ammonia-oxidizing bacteria (AOBs) and nitrite-oxidizing bacteria (NOBs) work cooperatively to transform ammonia into nitrate. Many of the waste products excreted by microbes still contain substantial chemical energy. This leads to a question: why aren’t these microbes further metabolizing their waste products to extract more ATP? In some cases, such as the cross-feeding between methanogens and aerobic methanotrophs, cross-feeding is strongly favored by environmental constraints: methanogenesis requires highly anaerobic conditions, while methanotrophy is typically an aerobic process (*10*). As a result, these processes must be separated. However, there are other cases where the reasons for cross-feeding are not clear. For example, in the case of nitrification, both ammonia and nitrite oxidation are aerobic processes, and could therefore occur within a single organism. Indeed, within the past decade, microbes performing both ammonia and nitrite oxidation, known as “comammox” bacteria, were discovered and isolated (*11*). Thus, the excretion of a compound as a waste product is not necessarily predetermined by chemistry, but may instead be contingent on ecological and evolutionary forces. Experimental evolution also supports this idea: a single strain of *Escherichia coli* in certain laboratory environments has evolved into a consortium that cross-feeds byproducts of sugar metabolism (*12*).

Thus, cross-feeding can be an outcome of an organism’s context — but what conditions promote cross-feeding? The first theoretical insights into waste-product cross-feeding emerged from the work of Heinrich and colleagues (*13, 14*). This body of work was concerned with calculating the optimization of metabolic pathways, and showed that optimal ATP flux in a static, nutrient-rich environment may be achieved at intermediate pathway lengths such that not all possible free energy is extracted from the initial substrate. This result suggests that cross-feeding might arise from organisms optimizing their own metabolism, creating byproduct pools that can then be capitalized on by other organisms. However, this analysis focused on the initial fitness of a new organism in an environment, which in general is not the same as fitness in an eco-evolutionary steady state.

Later theory by Doebeli showed that cross-feeding could evolve via adaptive dynamics, but in a phenomenological model that did not consider the intracellular state of the cells (*15*). A landmark work by Pfieffer and Bonhoeffer (PB) extended Heinrich’s model of multienzyme pathways to evolution in a chemostat, demonstrating that a cross-feeding consortium can be evolutionarily stable at high nutrient fluxes (*16*). PB argued that stable cross-feeding evolution depends on three factors: (1) ATP flux maximization, (2) minimization of the concentration of pathway enzymes, and (3) minimization of the concentration of pathway intermediates, justified as a form of metabolite ‘toxicity’. In their metabolic model, these factors were encoded as a linear penalty on ATP flux with increasing enzyme and intermediate concentrations. However, whether these principles translate beyond their particular model assumptions remains unclear. There are many possible forms of metabolite toxicity, do they all support cross-feeding? Additionally, PB did not differentiate between the evolution of abbreviated pathways in static environments and the evolutionary stability of cross-feeding. Does the evolution of an abbreviated pathway always lead to an evolutionarily stable cross-feeding consortium?

In this work, we develop a mathematically rigorous theory of the evolution of waste-product cross-feeding in a chemostat setting. We begin with the simplest possible metabolic model in which cross-feeding might arise. This model features a two-step pathway in which thermodynamics plays the role of toxicity. Surprisingly, we show that this model, despite favoring the evolution of abbreviated pathways and leading to the coexistence of distinct metabolic strategies, cannot support evolutionarily stable cross-feeding. We then explore an extended model that includes an osmotic-like metabolite penalty, and show that this penalty allows for evolutionarily stable cross-feeding. The key to cross-feeding stability is the ability of consortia to divide the burden of toxic metabolites among a larger population. Interestingly, we find that the transition to cross-feeding exhibits non-trivial dynamics, with community biomass and internal cellular contents changing discontinuously. Like PB, we find this transition occurs at high nutrient fluxes. Together, our analysis of these minimal models provides new insights into the requirements for evolutionarily stable cross-feeding, and highlights environmental and experimental conditions conducive to this outcome.

## Results

### Eco-evolutionary model

We first examine the simplest possible microbial metabolic model in which cross-feeding might be expected to arise. In this model, the unavoidable presence of thermodynamic product-feedback inhibition, i.e. the fact that all chemical reactions are in principle reversible, implies a form of metabolite toxicity. We consider the dynamics of strains of microbes with population density *ρ*_σ_ that exist within a chemostat with dilution rate *D*. We will consider multiple coexisting strains, but for simplicity suppress the strain index *σ*. The ratio of individual cell volume to chemostat volume is *r*_V_. A schematic of the model is shown in Fig. 1A. The microbes can extract energy from two chemical reactions: the transformation of compound 0 into compound 1, and the transformation of compound 1 into compound 2. This formalism can naturally be extended to an arbitrary number of reactions, but we consider this two-reaction system as a minimal model. Intracellular concentrations of these compounds are denoted *c*_*i*_, while extracellular concentrations are denoted *c*_*i,e*_ and external input fluxes denoted by *s*_*i*_. The transformation of *c*_*i−1*_ into *c*_*i*_ is catalyzed by enzyme *E*_*i*_ (for *i* = 1, 2) and the transport of *c*_*i*_ into and out of cells is performed by transporter *T*_*i*_ (for *i* = 0, 1, 2). We assume transport occurs passively. The total level of enzymes within a cell (including transporters) is constrained by the cell’s enzyme budget, here normalized to 1. The catalysis of *c*_*i*_−1 into *c*_*i*_ is limited by product inhibition, with equilibrium constant *K*_eq,*i*_. Each transporter has a rate parameter *β*_*i*_, which defines its performance relative to the catalytic enzymes. For brevity, we define catalysis and transport fluxes *J*_*c,i*_ and *J*_*t,i*_. The growth rate of a cell, *g*, is the sum of the cell’s catalysis fluxes, normalized by yield parameters *α*_*i*_ and reduced by the required energy flux for cell maintenance *g*_m_. To account for cell expansion, intracellular compounds are diluted at a rate proportional to *g*.

**Figure 1.**
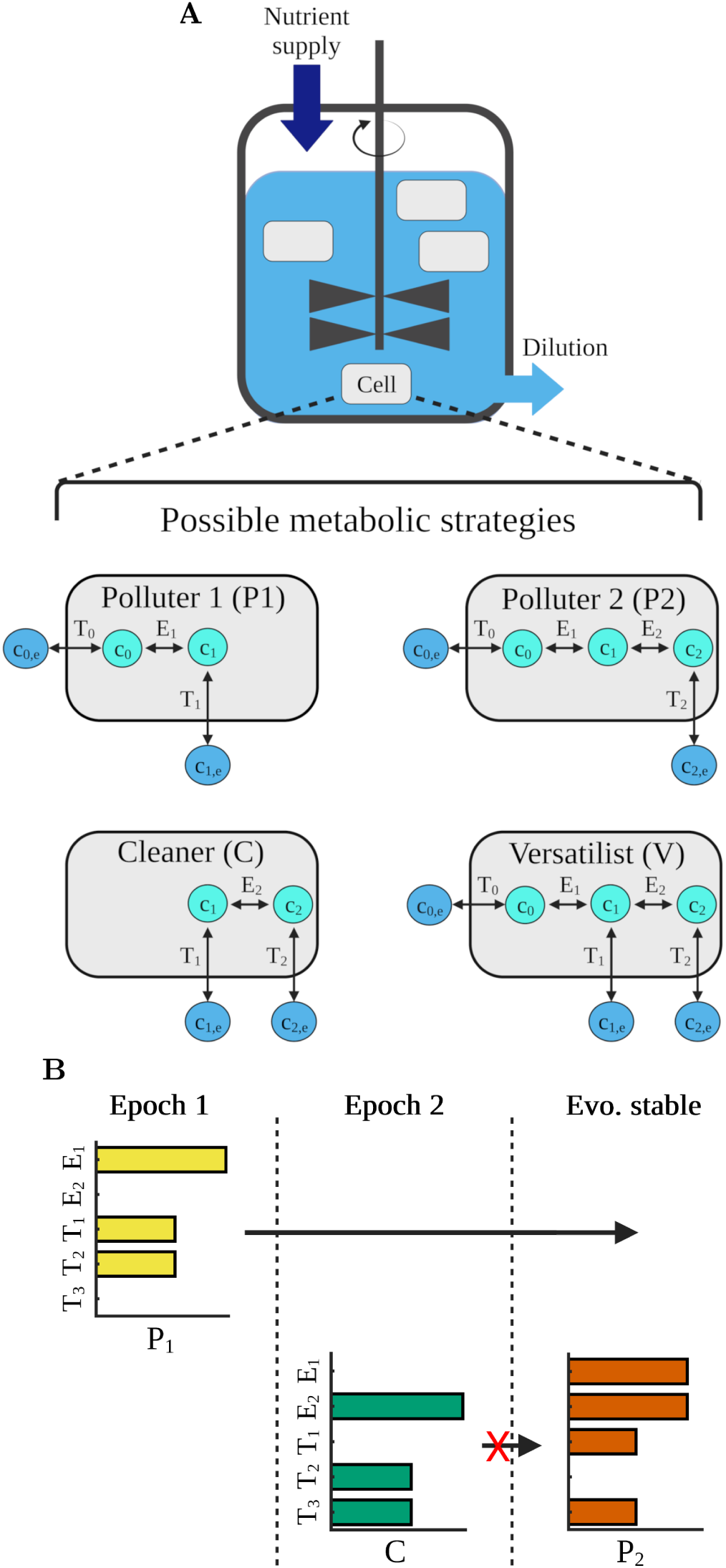
Eco-evolutionary model of microbial metabolism. (*A*) Elements of model: cells exist within a well-mixed vessel with constant dilution and nutrient addition; cells may allocate their budget between five different enzymes, gaining energy from up to two different reaction steps. (*B*) Representative example of an evolutionary algorithm trajectory leading to P1+P2 community: the ecosystem is first colonized by a P1 strain, and is then invaded by a C strain. The C strain is then displaced by an invading P2 strain. The resulting P1+P2 consortium is evolutionarily stable. Bars represent the logarithm of the strain’s investment in each enzyme.

The dynamics of the model are thus defined by the following equations:

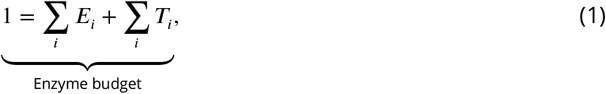

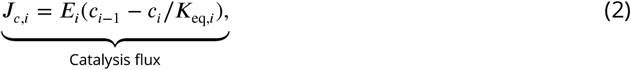

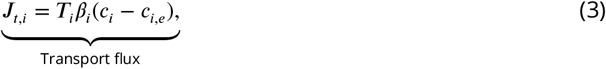

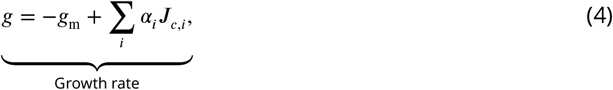

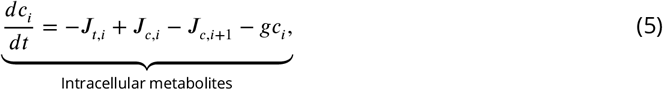

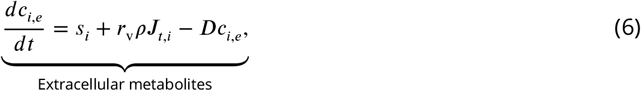

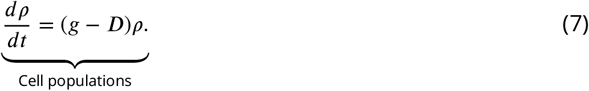

For simplicity, we initially set *K*_eq,*i*_ = *K*_eq_, *β*_*i*_ = *β*, and *α*_*i*_ = *α*. The equations shown here are the nondimensionalized version of the model, with the original dimensional model found in *SI Appendix* 1.

Before analyzing the model, we define the possible metabolic strategies that could arise. A cell could express only enzymes needed for the first reaction (*T*_0_, *T*_1_, and *E*_1_), and we refer to such a cell as a ‘Polluter 1’ (P1), since it pollutes the environment with compound 1. A cell could express only enzymes needed to perform both reactions (*T*_0_, *T*_2_, *E*_1_, and *E*_2_) and we refer to this cell as a ‘Polluter 2’ (P2). We refer to a cell expressing only the enzymes needed for the second reaction (*T*_1_, *T*_2_, and *E*_2_) as a ‘Cleaner’ (C) as it removes compound 1 from the system. In principle, cells could express all enzymes at once, and we refer to a cell of this type as a ‘Versatilist’ (V), but we do not find any conditions where this strain is evolutionarily stable in our models. Cross-feeding in our model would manifest as a stable consortium of P1 and C.

We are interested in the evolutionarily stable steady states of the above model, which we define as communities that cannot be successfully invaded by any other metabolic strategy. Note that this definition is agnostic to actual mechanisms of evolution (e.g. mutation), assuming that over long enough timescales, a stable state will eventually be reached. In the chemostat context, the evolutionarily stable state corresponds to a consortium that is able to generate an environment that no other cells can invade. We note the evident relation between this condition and the ‘pessimization principle’ of adaptive dynamics, in which invasion is prevented by a worsening of the environment (*17*). To find these stable states in our model, we employ a numerical evolution algorithm. We begin with an abiotic environment at steady state, and then compute the strain (i.e. metabolic strategy) with the highest possible invasion fitness in this environment. This strain is inserted into the system and allowed to reach steady state. We then compute the strain with the highest invasion fitness in the new environment, and this new strain is inserted. The above steps are repeated until no new strategy can invade. In Fig. 1B, we show an exemplary evolutionary trajectory. The initial abiotic environment is first invaded by a P1 strain, followed by invasion by a C strain. Finally, this C strain is driven to extinction by an invading P2 strain, resulting in an evolutionarily stable P1+P2 consortium.

### Evolutionarily stable states with thermodynamic toxicity

In Fig. 2A, we show the evolutionarily stable states of the thermodynamic toxicity model as a function of dilution rate, *D* and nutrient supply rate *s*_0_ at *K*_eq_ = 1. We find the final states are unique, independent of how we initialize the evolutionary algorithm. The corresponding population densities and extracellular compound concentrations as a function of dilution rate, *D* for an intermediate value of *s*_0_ are shown in Fig. 2C-D. At low values of supply rate and high values of dilution rate, the concentration of compound 0 is too low to allow persistence of any strain, regardless of strategy. In this regime, the extracellular concentration of compound 0 is simply *s*_0_/*D*. Eventually with increasing supply rate (or decreasing dilution rate), P1 becomes able to invade and persist in the chemostat. Both the population density of P1 and the extracellular concentrations of compounds 0 and 1 increase with decreasing dilution rate. The community remains exclusively composed of P1 until the supply rate increases sufficiently (or dilution rate decreases sufficiently), at which point a two-strain consortium forms. Interestingly, this consortium is not a cross-feeding pair, but rather is composed of P1 and P2. These two strategies both feed on compound 1 but excrete different byproducts. This transition between P1 and a P1+P2 consortium occurs once P1 has polluted its environment with enough compound 1 to substantially product-inhibit the reaction carried out by enzyme 1. Once this occurs, P2, which is not inhibited by extracellular compound 1 (because it lacks transporter T1), can successfully invade. Further increases in the supply rate or decreases in the dilution rate do not alter the strain composition of this consortium, with the P1+C cross-feeding consortium never being evolutionarily stable. The ratio of P1 and P2 is variable, with each strain increasing at different rates with decreasing dilution rate. The P1 to P2 ratio also varies as a function of supply rate, though the individual populations increase nonlinearly with increasing supply. Extracellular concentrations of all compounds increase with decreasing dilution and increasing supply rate in the P1+P2 regime.

**Figure 2.**
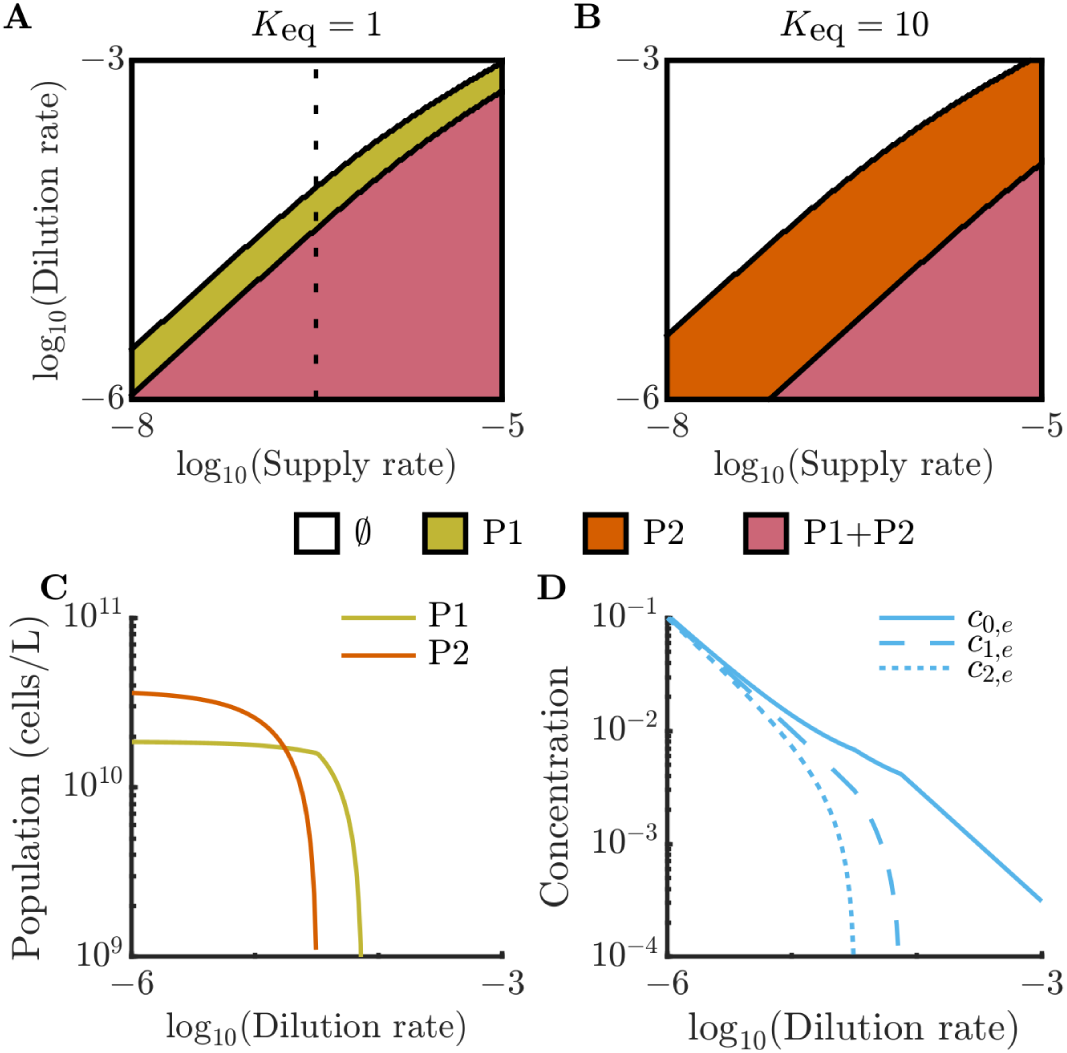
Evolutionarily stable communities in the thermodynamic toxicity model. (*A*) Phase diagram of evolutionarily stable community states with *K*_eq_ = 1 as a function of nutrient supply rate, *s*_0_, and dilution rate, *D*. Model parameters can be found in *SI Appendix* 1. “∅” indicates environments where no strains can persist. As in *A*, but with with *K*_eq_ = 10. (*C*) Population densities of P1 and P2 and (*D*) extracellular compound concentrations in evolutionarily stable communities as a function of dilution rate *D* along dashed line of communities seen in panel *A*.

What occurs if the strength of product inhibition is altered? In Fig. 2B we show the equivalent phase diagram for a system with *K*_eq_ = 10. In this case, product inhibition is weakened such that P2, which performs both reaction steps, is the first strategy that can invade the environment. However, a large enough increase in supply rate or decrease in dilution rate lead to P2 polluting its own environment with external compound 2, allowing invasion by P1 and leading again to a P1+P2 consortium. Thus, while the thermodynamic toxicity model readily supports the evolution of an abbreviated metabolic pathway strategy P1, this does not necessarily lead to cross-feeding, instead we find coexistence of two different types of polluter, P1 and P2.

We have thus far considered two energy-yielding reactions that provide the same energy yield, but what occurs when these two yields are substantially different, as is the case in nitrification (*11*)? Surprisingly, we find that even if the second reaction yields no energy (i.e. *α*_2_ = 0), the P1+P2 consortium still arises. This is due to the fact that the ability of P2 to invade a P1 community is not simply due to the additional energy yield from the second reaction, but rather to the fact that P2 is not inhibited by the metabolic byproduct of P1 (Fig. S1). Similarly, we find that varying the *K*_eq,*i*_ between the two reactions results in similar evolutionarily stable consortia (Fig. S2).

Can a cross-feeding consortium ever be stable in the thermodynamic toxicity model? Extensive numerical exploration of the model identified no conditions under which cross-feeding is stable, strongly suggesting the model cannot support this behavior. In addition to numerical analysis, we analytically characterized the evolutionary steady states of a slightly simplified, more tractable version of the thermodynamic toxicity model that exhibits near-identical evolutionarily stable states. In this simplified version of the model, the intracellular dilution term *gc*_*i*_ in Eq. 5 is neglected. In *SI Appendix* 5, we show that this version of the model does not permit stable cross-feeding. In particular, we show that the environment produced by the optimal P1+C consortium is always invadable by the optimal P1+P2 consortium. From these numerical and analytical results, we conclude that thermodynamic toxicity alone is not sufficient to produce evolutionarily stable cross-feeding. Note that this result challenges predictions of previous work on cross-feeding evolution which assumed that a stable cross-feeding relationship will necessarily emerge if the initial optimal invader has an abbreviated metabolic pathway (*13, 14*).

To intuitively understand why thermodynamic toxicity does not support cross-feeding, one can conceptualize a P1+C consortium as effectively a P2 cell split into two compartments, as shown in Fig. 3A. In net, P1+C are performing the same reactions as P2, but the P1+C consortium must use a portion of its enzyme budget on *T*_1_ transporters that P2 does not require. This inefficiency makes the P1+C consortium always underperform relative to the best P2 strain, meaning that any P1+C ecosystem is susceptible to invasion by P2. This difference in efficiency can be seen in the steady-state extracellular concentrations *c*_0,*e*_ of optimal P1+C and P1+P2 communities grown under the same conditions: evolutionary stability within a chemostat is dependent on strains’ ability to create and survive the harshest possible environment. Thus, the evolutionarily stable community will be able to deplete nutrient in the environment to the lowest possible level. In Fig. 3B, we show the ratio between the steady-state *c*_0,*e*_ concentration reached by the optimal P1+C community to that reached by the optimal P1+P2 community. As can be seen, this ratio is always greater than 1, becoming closer to 1 at larger nutrient supply rates, and becoming larger when thermodynamic toxicity is weak (i.e., large *K*_eq_). The inefficiency of the cross-feeding pair is also reflected in the intracellular concentrations within the consortia. In Fig. 3C, we show the internal concentrations of *c*_0_ and *c*_2_ in optimal P1+C and P1+P2 communities growing under the same conditions, normalized to the abiotic external steady-state concentration *c*_0,*e*_ in the absence of growth. As can be seen, the P1+P2 community can operate at the same growth rate while sustaining a lower intracellular concentration of its substrate *c*_0_. As supply rate grows, the relative difference in intracellular concentration of *c*_0_ and the toxic end product *c*_2_ decreases. This occurs because the absolute flux of substrate is increasing, such that a smaller relative concentration difference is needed to support the required growth rate *D*. A version of Fig. 3 generated from our simplified model neglecting intracellular dilution is shown in Fig. S4 and shows similar trends.

**Figure 3.**
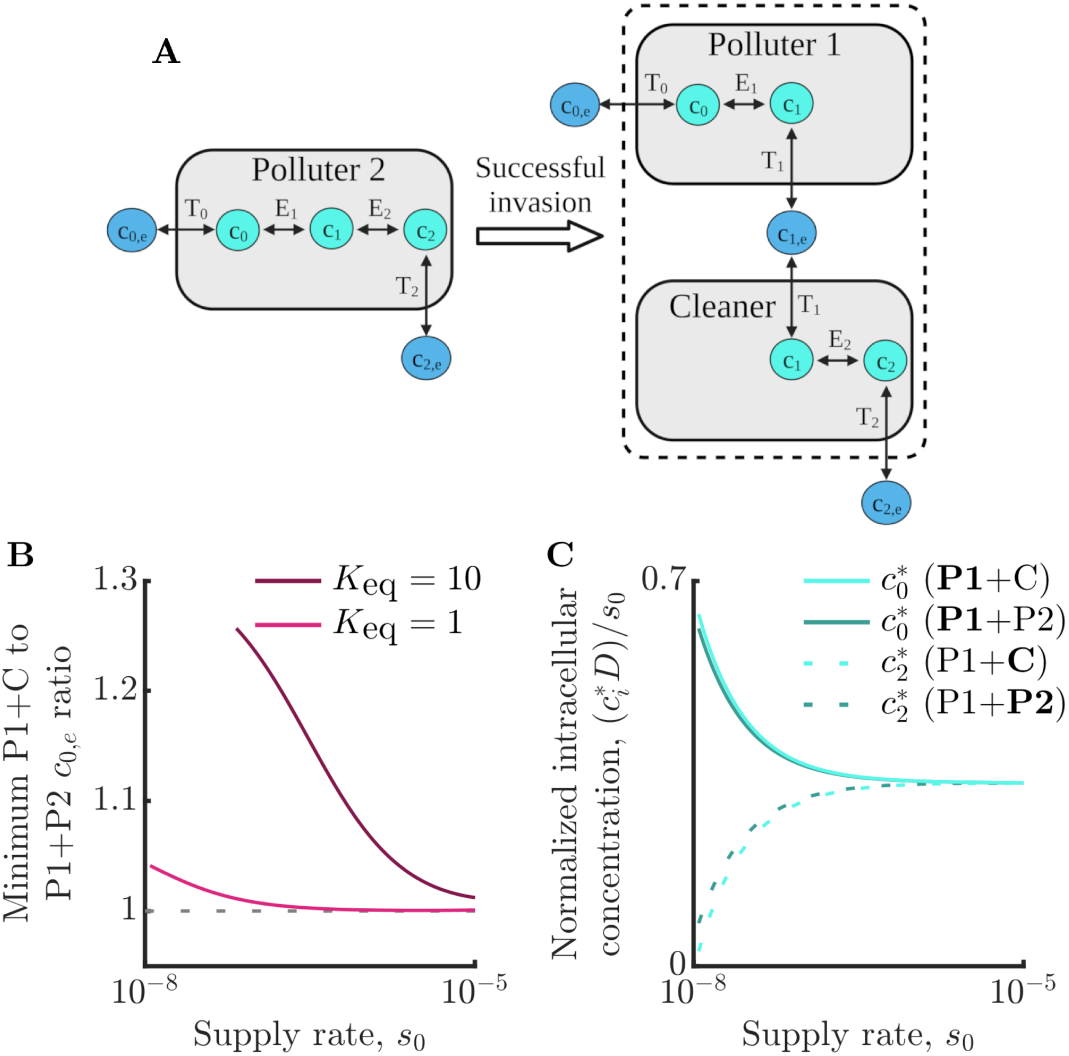
Cross-feeding is never evolutionarily stable in the thermodynamic toxicity model. (*A*) Schematic of P2 invasion of a P1+C cross-feeding community. The P1+C consortium can be conceptualized as a P2 cell split across two cells. Due to the P1+C community’s need for additional transporters, it is always invadable by P2 in the thermodynamic toxicity model. (*B*) Ratio of the steady-state extracellular concentration of compound 0, *c*_0,*e*_, generated by the optimal P1+C and P1+P2 consortia. Dashed line indicates a ratio of one. A ratio of greater than one indicates that the optimal P1+C community can be invaded by a P2 strain. (*C*) Normalized steady-state intracellular concentrations *c*_*i*_ of compounds 0 and 2 in optimal P1+C and P1+P2 communities as a function of supply rate *s*_0_. Concentrations are normalized by *s*_0_/*D*, the steady-state concentration of *c*_0,*e*_ under abiotic conditions, to better visualize differences between consortia. In both P1+C and P1+P2 consortia, 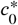 is the concentration within P1 cells, while 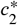 is the concentration within C cells in the P1+C consortia and within P2 cells in the P1+P2 consortia. For panels *B* and *C, D* = 10^−6^, and all other model parameters can be found in *SI Appendix* 1.

### Nonlinear toxicity enables stable cross-feeding

Cross-feeding is evolutionarily *unstable* in the minimal thermodynamic model we have explored thus far, but is widespread in natural microbial communities. What might favor cross-feeding in nature? Based on the above analysis, cross-feeding will be favored if splitting up a metabolic pathway between two cells is beneficial. This would follow from an additional penalty on internal metabolites beyond product-feedback inhibition. Thus, we now explore a version of the model with an additional, nonlinear metabolite-toxicity penalty. Specifically, we introduce a toxicity Ω that modifies the growth rate such that 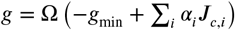 reduces the growth rate in proportion to the total intracellular compound concentration: Ω = 1 − ∑_*i*_*c*_*i*_. We refer to this penalty as “osmotic-like” as it places a limit on the total intracellular compound concentration, analogous to an osmolarity constraint in a given extracellular environment. The primary difference between this penalty and the thermodynamic penalty is that, while for the thermodynamic penalty increasing all intracellular compound concentrations in proportion always led to a growth rate increase, the dependence on compound concentration is now nonmonotonic, as depicted in Fig. 4A.

**Figure 4.**
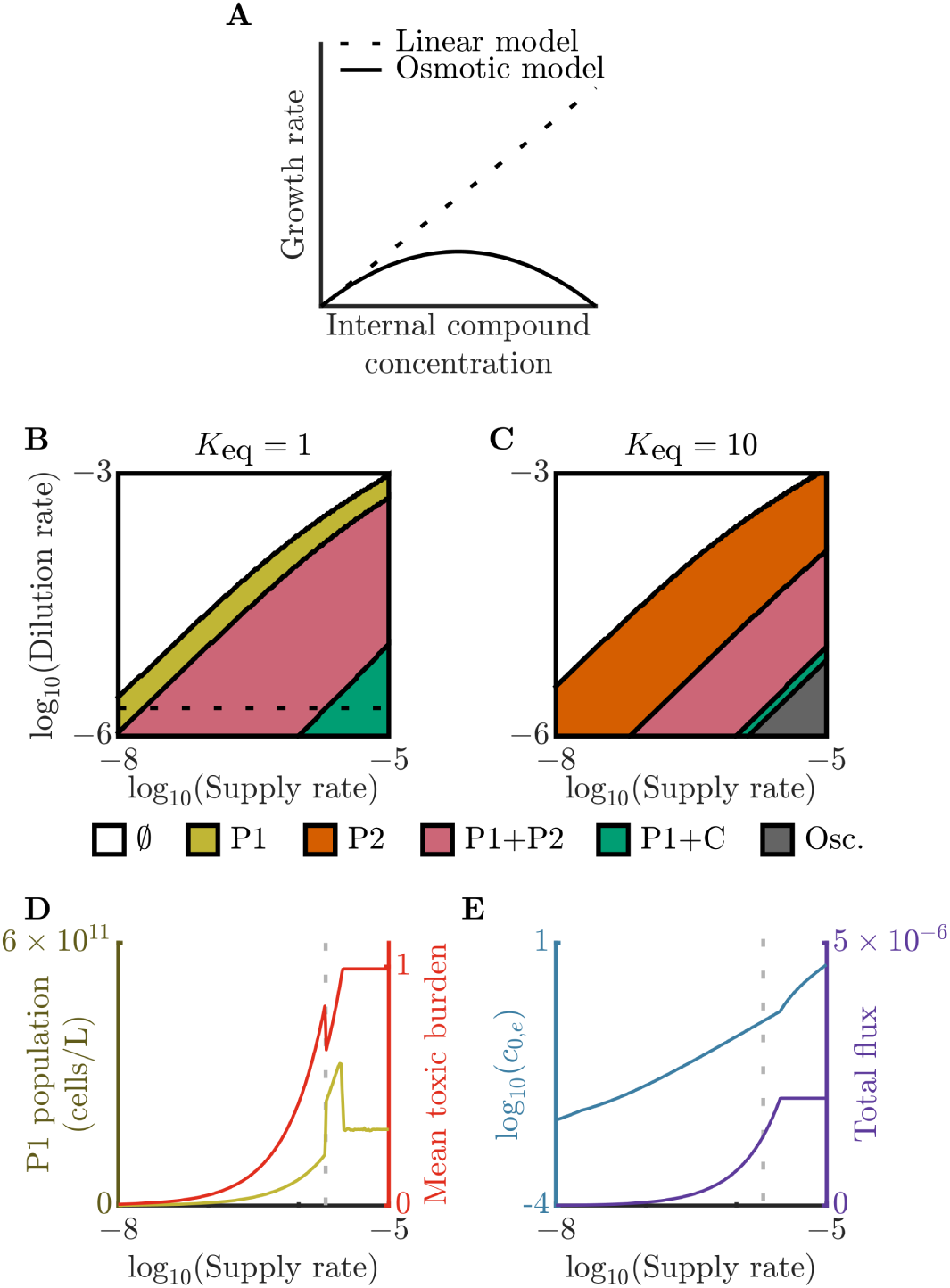
Osmotic toxicity allows for evolutionarily stable cross-feeding. (*A*) Example of growth rate as a function of internal compound concentration for linear thermodynamic model and nonlinear osmotic-penalty model. Phase diagrams of evolutionarily stable communities as functions of nutrient supply rate, *s*_0_, and dilution rate, *D*, for (*B*) *K*_eq_ = 1 and (*C*) *K*_eq_ = 10. Model parameters same as in Fig. 2*A*-*B*. “Osc.” denotes oscillatory states for which evolutionarily stable communities cannot be computed using our framework. (*D*) P1 population density and mean community toxic burden, (*E*) extracellular compound 0 concentration and total flux, all as functions of supply rate, *s*_0_, along the dashed line in *B*. Mean toxic burden is the sum of intracellular metabolite concentrations ∑_*i*_*c*_*i*_ averaged over the community. Total flux is defined as the flux of compound 0 that is consumed by the evolutionarily stable consortium. The dashed line denotes the point at which the cross-feeding P1+C community becomes evolutionarily stable.

What are the evolutionarily stable states with an osmotic-like toxicity? In Fig. 4B-C We show the phase diagram of evolutionarily steady states for *K*_eq_ = 1 and *K*_eq_ = 10, analogous to Fig. 2A and B. At high values of dilution rate and low values of supply rate, the model behaves much like the thermodynamic model: with decreasing dilution and increasing supply, the system transitions from abiotic to supporting a strain (P1 or P2), before eventually leading to a consortium of P1+P2. However, unlike the original model, further increasing the supply rate or decreasing dilution rate leads to a stable cross-feeding P1+C consortium. Why is cross-feeding now stable? To understand this, we must consider the total compound load in the chemostat. In the absence of microbes, the steady-state level of *c*_0,*e*_ in the system is *s*_0_/*D*. Thus microbes growing at higher supply rates and lower dilution rates deal with an increasingly high ambient level of substrate. In the osmotic-like toxicity model, substrate level now influences the growth rate in a nonmonotonic manner, and increasing the substrate influx can be detrimental to the cells. Thus, in this high-toxicity regime, it becomes beneficial to split metabolism between two bacterial populations that each individually will have lower toxic burdens. This can be seen in Fig. 4D: at the cross-feeding transition point, the number of cells in the population jumps discontinuously while the mean toxic burden of the cells drops discontinuously. Interestingly, these discontinuities in community properties do not extend into the environmental properties. In Fig. 4E we show the steady-state value of substrate flux and of *c*_0,*e*_, both of which change continuously across the transition.

We find that within the cross-feeding regime there is a second transition as supply rate increases further, corresponding to a plateau in the community’s metabolic flux. At this transition, we see a rapid but continuous change in community biomass and enzyme strategies. The biomass rapidly drops, coinciding with a shift in both the P1 and C enzyme strategies, lowering the level of their influx transporter (e.g., *T*_0_ in P1) and increasing the levels of their metabolic enzyme and outflux transporter (see Fig. S8). This transition represents the onset of ‘pessimization’ (*17*), a phenomenon in which a population severely degrades its environment to repel competitors. Indeed, a P1 strain evolved at a supply rate just before the onset of pessimization cannot survive in the post-transition condition due to a much higher internal toxic burden. This abrupt pessimization of the environment is limited by the nonmonotonic growth function: a cell can only support a particular maximum flux while still growing at the dilution rate. This limit leads to the plateau in metabolic flux, mean toxic burden, and community biomass (Fig. S8).

Note that in addition to the cross-feeding regime, further decreases in dilution rate and increases in supply lead to a regime with oscillatory ecological dynamics. These oscillations occur due to cycles of environmental toxification and detoxification. In this manuscript, we do not attempt to identify the evolutionarily stable consortia within this regime. Our model framework neglects dynamic metabolic regulation, which is a valid assumption for environments that reach time-invariant steady states, such as those analyzed above. In a dynamically varying environment, regulation becomes critical (*18*) and thus a more complex model formulation would be required. In addition to the oscillatory regime, the osmotic toxicity model can also exhibit bistability in invasion outcome when the initial population of an invading strain is small. We examine this phenomenon further in *SI Appendix* 6.

The form of nonlinear toxicity we consider in the osmotic model is strong, potentially driving cell growth rate to zero in severe cases. Will cross-feeding occur with a weaker form of nonlinear inhibition? To explore this, in *SI Appendix* 7, we examine the evolutionary steady states of a model with saturating (Monod-like) growth kinetics. In this case, the inhibition causes the growth rate to plateau at high intracellular concentrations rather than go to zero. We find no cross-feeding in this case in the parameter regime explored in Fig. 4BC, but we do not completely rule out the possibility of cross-feeding, particularly in parameter regimes with extremely high supply rates and low dilution rates.

## Discussion

In this work, we develop a minimal evolutionary theory for the emergence of microbial cross-feeding. We first explore a model in which metabolism is constrained by a limited enzyme budget and the thermodynamic reversibility of chemical reactions. Contrary to predictions of early cross-feeding theory, this model does not support evolutionarily stable cross-feeding, despite the emergence of abbreviated metabolic pathways. Rather than a cross-feeding community, a consortium emerges of microbes consuming the same nutrient but secreting different waste products. When a nonlinear, osmotic-like constraint on internal metabolite levels is added to the model, a cross-feeding consortium emerges in environments with high nutrient flux and low dilution rate. Intuitively, this occurs because once the penalty for carrying intracellular metabolites is superlinear in metabolite concentrations, it becomes beneficial to split metabolism between two cells. The average penalty for two populations, each with a smaller individual metabolite pool, is less severe than that for one population with a single, larger pool. The emergence of cross-feeding is likely not unique to the particular osmotic-like penalty we have explored here — we expect that any sufficiently nonlinear penalty would enable cross-feeding.

While our model focuses on evolutionary stability, its conclusions can also be applied to the prediction of ecological community states. In an ecosystem that is invaded by many variants of P1, P2, and C strains, the ecologically stable consortium will approximate the evolutionary stable consortium we find in our model. Thus, we predict that cross-feeding consortia will dominate in high nutrient flux environments, while low-flux environments will feature a single pathway length (either abbreviated or long), and intermediate fluxes will promote coexistence of multiple pathway lengths beginning at the same nutrient. Note that despite the differences between our evolutionary theory and the earlier optimization theory of Heinrich and colleagues, the prediction that crossfeeding only occurs at high nutrient fluxes is shared between the two theories. In terms of empirical support, Kreft et al. recently argued that the prediction that cross-feeding is favored at high flux is consistent with the observed difference in cross-feeding prevalence between anaerobic and aerobic ecosystems (*19*). The authors’ argument is based on the fact that cross-feeding consortia tend to be more common in anaerobic conditions than aerobic conditions, and that these anaerobic environments are anaerobic precisely because they receive high nutrient fluxes (specifically that the high carbon flux creates a large oxygen demand which depletes oxygen). In future, such empirical support could be strengthened by quantitative comparisons between environmental parameters and local microbial community composition. Recent work by Zhang et al. (*20*) provides a potential basis for such analyses. The authors use metagenomically-assembled genomes (MAGs) to assess splitting of nitrification pathways in marine microbes across multiple geographic locations and timepoints, finding many organisms with only partial nitrification pathways. Correlating such information with data on local substrate fluxes would enable testing of theoretical predictions. In addition to changes in community composition, our model also predicts the existence of a discontinuity of total community biomass as nutrient flux increases. While in principle detectable empirically, given the relatively small size of the discontinuity with our parameterization and the noisiness of environmental data, such a signal may not be easily detectable in data.

Here we explored one model architecture that permits cross-feeding, but how might the predicted relationship between evolutionary steady states and environmental conditions vary under different model structures and assumptions? To assess this, we compare our model to the theory of Pfieffer and Bonhoeffer discussed earlier (*16*), as their model has a somewhat similar structure and also permits cross-feeding. We focus this comparison on dilution rate, as this is an environmental parameter explored in both our work and the PB model. The fact that PB’s model supports cross-feeding is consistent with our results, given that the growth penalty PB impose on intracellular metabolites is strongly nonlinear. The overall community diversity is also similar between the models: the evolutionary stable states contain at most two strains. At high dilution rates, the predicted evolutionarily stable states of both models are also similar: no strains survive when dilution is too high, and the first communities to emerge when dilution is lowered are composed of single strains that either entirely or partially degrade the supplied substrate. However, the models disagree on behavior at intermediate dilution rates. In our model, as dilution rate further decreases the P1+P2 consortium becomes stable, and is then succeeded by the P1+C consortium which is stable until the ecosystem destabilizes into an oscillatory state. By contrast, in the PB model, the equivalent of the P1+C consortia becomes stable first, and is then succeeded by the P1+P2 consortia at lower dilution rate, which itself then gives way to a P2 community at even lower dilution rate. Thus, while both models predict cross-feeding to be stable only at lower dilution rates, the exact structure of the phase diagrams differs significantly. Therefore, there is no universal phase diagram among evolutionary models that permit cross-feeding.

We have primarily compared our theory to other very general theories of pathway splitting and cross-feeding emergence, but there is also a relevant body of theory focusing on one particular class of pathway splitting, namely overflow metabolism. In overflow metabolism, known in other contexts as the Crabtree or Warburg effect, organisms partially utilize a shorter, lower yield fermentative pathway in addition to an oxidative one despite having access to oxygen. This behavior occurs across the tree of life, from bacteria to single-celled eukaryotes to mammalian cancer cells (*21*–*23*). For example, at high growth rates *E. coli* metabolizes glucose into acetate and excretes some of this product rather than further respiring it. Recent elegant theoretical work by de Groot et al. shows that many existing theories of overflow metabolism can be unified into a single framework in which the optimality of overflow metabolism requires two growth-limiting constraints (*24*). Our work adds further depth to this picture by considering evolutionary dynamics. We show that, like ecological optimality, the evolutionary stability of abbreviated pathways can occur with only two constraints (in our case, arising from an enzyme budget and thermodynamic toxicity). However, we show that not all constraints are equal when it comes to community composition: cross-feeding consortium stability depends on the specific constraints. Thus, the simple two-constraint theory of de Groot et al. is not fully applicable to an evolutionary context. Note, however, that there is a fundamental difference between the abbreviated pathway utilization which arises in our model and that which occurs with overflow metabolism. Our model does not support evolutionary stability of combined pathway utilization, which commonly occurs in models of overflow metabolism. Indeed, the mixed pathway utilizing versatilist is never evolutionary stable in our model.

In addition to providing an informative comparison, the theoretical literature on overflow metabolism also provides a guide for the future study of evolutionary cross-feeding theory. A promising direction of further research would be to elucidate more rigorously the constraint properties that enable cross-feeding evolutionary stability. We hypothesize this constraint property is non-monotonicity, as a Monod growth function (which is concave but monotonic, see *SI Appendix 7*) did not produce cross-feeding, but the non-monotonic osmotic growth function did. With such an understanding, one could attempt to unify models such as ours and that of PB, as was done for disparate models of overflow metabolism by de Groot et al.(*24*). Another fruitful direction would be to explore the non-steady-state regimes we find in our model. At very high values of nutrient supply, the ecosystem experiences periodic oscillations of pollution and detoxification. Our current framework does not allow for determination of possible evolutionary steady states in such conditions, not only due to the manner in which we compute invasion fitness, but also due to the importance of enzyme regulation in such fluctuating environments. Additionally, fluctuating nutrient environments could promote metabolic division of labor as such communities could adjust their metabolic activities by simply adjusting their population abundances. To develop a framework for tackling such issues, recent work on adaptive dynamics and enzyme regulation in fluctuating environments (*18, 25*) may be a useful starting point.

Our primary focus in this work was the stability of metabolic cooperation, but in analyzing our models we also identified more competitive types of metabolic interaction based on environmental pollution. In both the thermodynamic and osmotic versions of our model, we found two-strain consortia that coexist on a single limiting resource despite not engaging in cross-feeding. Instead, these organisms establish niches based on how they pollute their environment, which each strain emitting a unique pollutant. While not additional resources, these pollutants are additional ‘limiting factors’ that in accordance with the generalized competitive exclusion principle allow multi-strain coexistence (*26*). This behavior is similar to that identified by Großkopf et al. (*27*). Parallels can be drawn between the different polluter strains in our model and the multiple different types of waste products that emerge from incomplete utilization of real-world substrates, e.g. different forms of sugar fermentation resulting in acetate, lactate, or ethanol. Our model suggests that this diversity of pathways may emerge as a result of niche creation: a strain emitting a novel pollutant can persist even if that pathway has inferior energetic or kinetic properties. In addition to the evolution of pollutant diversity, our model also exhibits regimes of severe ‘pessimization’ (*17*) in which the evolutionarily stable community severely degrades its environment. This is seen most strikingly in Fig. 4D and Fig. S8, which displays a region of parameter space in which the steady-state biomass of the cross-feeding consortia decreases despite the nutrient supply increasing. This highlights another counter-intuitive benefit of polluting: even if self-pollution severely damages the resident community, polluting may still be evolutionarily favored if it can prevent the growth of potential competitors.

We have framed our theory around bacterial cross-feeding evolution, but given the simplicity of the framework, our results may be useful in explaining metabolic cooperation among single-celled organisms across the tree of life. Generally, our results suggest that given the many strong nonlinearities inherent in metabolism, it is not surprising that cooperative metabolic cross-feeding is so common in the natural world.

## Methods

Details on model construction, parameterization, and the analysis methods employed can be found in the *SI Appendix*. All codes and simulation results can be found at https://github.com/jamie-alc-lopez/waste_crossfeeding_evolution.

## Acknowledgments

We thank the Donia lab, Wingreen lab, Orkun Soyer, Xin Sun, and Jean Vila for productive discussions. JAL was supported by a Stanford Baker Fellowship. The work by Bo Liu was done when he was an undergraduate in Integrated Science at Yuanpei College, Peking University. This research was supported by NIH grant R01 GM082938. This research was also supported in part by grants NSF PHY-1748958 and PHY-2309135 and the Gordon and Betty Moore Foundation Grant No. 2919.02 to the Kavli Institute for Theoretical Physics (KITP). This work was performed in part at Aspen Center for Physics, which is supported by National Science Foundation grant PHY-1607611.

