## Supplementary material for "When is microbial cross-feeding evolutionarily stable?": SI Appendix

[a]Lewis-Sigler Institute for Integrative Genomics, Princeton University, New Jersey 08544, USA; [b]Department of Bioengineering, Stanford University, Stanford, CA 94305, USA; [c]Department of Applied Physics, Stanford University, Stanford, CA 94305, USA; [d]Center for Brain Science and Department of Molecular and Cellular Biology, Harvard University, Cambridge, MA 02138, USA; [e]Kempner Institute for the Study of Natural and Artificial Intelligence, Harvard University, Cambridge, MA 02134, USA; [f]Center for Quantitative Biology, Academy for Advanced Interdisciplinary Studies, Peking University, Beijing, 100871, China; [g]Peking-Tsinghua Center for Life Sciences, Academy for Advanced Interdisciplinary Studies, Peking University, Beijing, 100871, China; [h]Department of Molecular Biology, Princeton University, New Jersey 08544, USA

**3 Thermodynamic toxicity model with zero energy yield from second metabolic reaction** **5**

**4 Thermodynamic toxicity model with varying $K_{\mathbf{eq},i}$** **5**

**5 Derivation of evolutionarily stable strategies in a simplified thermodynamic model** **5**

**6 Bistability in the osmotic toxicity model** **8**

**7 Thermodynamic toxicity model with Monod growth kinetics** **10**

**8 References** **11**

### 1. Description of chemostat model

**A. Dimensional model equations .** We begin by describing the fully dimensional version of the model. We model a well-mixed population of metabolically distinct strains growing in a chemostat, each with cell number density $\rho_\sigma(t)$. The ratio of the volume of an individual cell and the total chemostat volume is $r_{\rm V}$. Each of $n$ compounds is supplied to the chemostat at rate $\hat{s}_i$ (concentration per time) and both nutrients and cells are diluted at rate $\hat{D}$. The extracellular concentration of each compound is denoted by $\hat{c}_{i,e}$. We also model intracellular concentration of each compound, denoted by $\hat{c}_i$ (leaving off the subscript $\sigma$ for compactness). Each cell has an amino acid budget, $B$, which represents the total number of amino acids it can use for protein synthesis, a fraction $\xi$ of which are in metabolic enzymes. Each cell has up to $n$ types of transporter enzymes, which transport compounds across the cell membrane with transport coefficient $\hat{\beta}_i$. This transport is modeled as passive, with the driving force being the difference between intracellular and extracellular concentrations. Each cell has up to $n-1$ types of metabolic enzymes, the abundances of which are denoted by $\hat{E}_i$, that process compound $i-1$ into compound $i$ with rate coefficient $\hat{\nu}_i$. This processing is assumed to be reversible, with equilibrium constant $K_{{\rm eq},i}$. Note, however, that we prohibit the net processing rates from being negative to prevent unrealistic negative growth rates. Each molecule of compound $i-1$ converted to compound $i$ yields $\Upsilon_i$ molecules of ATP. This ATP flux contributes to the growth rate $\hat{g}$ with a conversion coefficient $\hat{\alpha}_i$. The cell also requires a minimum substrate flux for cell maintenance, which reduces the growth rate by $-\hat{g}_{\rm m}$. Growth increases the number density of cells, but also dilutes intracellular compound concentrations. When modeling osmotic stress and other forms of toxicity, the cell's growth is reduced in proportion to its total intracellular compound pool, with toxicity concentration scale $c^*$. The equations governing the dynamics are thus:

$$\xi B = \sum_i \hat{E}_i + \sum_i \hat{T}_i, \qquad \text{[S1]}$$

$$\hat{J}_{c,i} = \hat{E}_i \hat{\nu}_i (\hat{c}_{i-1} - \hat{c}_i / K_{{\rm eq},i}), \qquad \text{[S2]}$$

$$\hat{J}_{t,i} = \hat{T}_i \hat{\beta}_i (\hat{c}_i - \hat{c}_{i,e}), \qquad \text{[S3]}$$

$$\hat{\Omega} = 1 - \frac{\sum_i \hat{c}_i}{c^*}, \qquad \text{[S4]}$$

$$\hat{g} = \hat{\Omega}\left(-\hat{g}_{\rm m} + \sum_i \hat{\alpha}_i \Upsilon_i \hat{J}_{c,i}\right), \qquad \text{[S5]}$$

$$\frac{d\hat{c}_i}{d\tau} = -\hat{J}_{t,i} + \hat{J}_{c,i} - \hat{J}_{c,i+1} - \hat{g}\hat{c}_i, \qquad \text{[S6]}$$

$$\frac{d\hat{c}_{i,e}}{d\tau} = \hat{s}_i + r_{\rm v}\rho \hat{J}_{t,i} - \hat{D}\hat{c}_{i,e}, \qquad \text{[S7]}$$

$$\frac{d\rho_\sigma}{d\tau} = (\hat{g} - \hat{D})\rho_\sigma, \qquad \text{[S8]}$$

where the $\hat{J}_{c,i}$ are the intracellular catalysis fluxes, the $\hat{J}_{t,i}$ are the transport fluxes, and $\hat{\Omega}$ is the osmotic growth penalty.

**B. Model parameterization.** For our parameterization, we use *Escherichia coli* as a reference organism. Our goal is for the resulting model to be a linear approximation of the real growth and transport dynamics of *E. coli*. When possible, we take values from *E. coli* growing on glucose via oxidative fermentation, such that the per-glucose ATP production is $\Upsilon = 12$ (1). We assume a 1 L chemostat, such that $r_{\rm V} = 2 \times 10^{-15}$ with a cell volume of $\approx 2$ fL (2). We take $\hat{s}_i$, $\hat{D}$, and $K_{{\rm eq},i}$ as free parameters that we vary in our analyses. For simplicity, we now assume all transporters and enzymes are equally efficient ($\hat{\nu}_i = \hat{\nu}$ and $\hat{\beta}_i = \hat{\beta}$).

We first estimate the total amino-acid budget of the cell $B$. We estimate this value from (3), which characterized proteome allocation in *E. coli*. In particular, we begin with $9.5 \times 10^8$ amino acids per fL, the total number per volume of amino acids in the proteome of an *E. coli* growing in rich media at a growth rate of 0.82 h$^{-1}$ (see Table S1 of (3)). With a cell volume of 2 fL (2), this translates to $B = 1.9 \times 10^9$ amino acids per cell. Note, however, that cells do not devote their entire budget to metabolism-related enzymes. In (1), it was found that cells only devote approximately ten percent of their budget to these functions, and thus we take $\xi = 0.1$.

Next, we estimate $\hat{\alpha}_i$, the constant that converts intracellular ATP flux to a growth rate. We begin with the per-mole ATP yield of 13.9 grams dry weight of cell biomass measured in (4). We first convert this to the intracellular concentration of ATP required to make one cell, assuming a dry cell mass of 0.278 pg (5) and a cell volume of 2 fL (2), giving us 10 M ATP/cell. In order to achieve a growth rate of 1 h$^{-1}$, the cell must therefore sustain a flux of 10 M ATP/h, giving us $\hat{\alpha} = 0.1$ (M ATP)$^{-1}$.

We now estimate the maintenance growth rate term, $\hat{g}_{\rm m}$. We decompose this term into three components: $\hat{g}_{\rm m} = \hat{\alpha}_{\rm m}\Upsilon_{\rm m}\hat{J}_{\rm m}$, where the three factors respectively represent the ATP-to-growth conversion, ATP yield per substrate, and substrate flux associated with maintenance. The maintenance substrate flux of *E. coli* growing aerobically on glucose is 1 mmol substrate per gram dry weight per hour (6). Given the dry cell mass and cell volumes used above, this gives $\hat{J}_{\rm m} = 0.14$ M glucose/h. Given

The authors declare no conflict of interest.

[1] JAL contributed equally to this work with BL.

that glucose utilization under anaerobic conditions generates $\Upsilon_{\mathrm{m}} = O(10^1)$ ATP (1) and assuming $\hat{\alpha}_{\mathrm{m}} = \hat{\alpha} = 0.1$ (M ATP)$^{-1}$ as calculated above yields a maintenance growth rate of $g_{\mathrm{m}} \approx 0.1$ h$^{-1}$

We next estimate $\hat{\nu}_i$, the constant relating the intracellular metabolite concentrations and the fraction of enzyme budget spent on metabolic enzymes to metabolic flux. We assume all $\hat{\nu}_i$ are equal such that $\hat{\nu}_i \equiv \hat{\nu}$. We begin with a measurement of ATP flux per proteome fraction devoted to fermentation from (1). This measurement, taken from Figure 4 of the reference, is 0.75 M ATP per $\mathrm{A}_{600\mathrm{nm}}$ per hour per protein fraction, where $\mathrm{A}_{600\mathrm{nm}}$ is the optical density of *E. coli* culture. Approximately one-tenth of the fermentation enzymes used to compute this measurement are transport enzymes, and thus we adjust this measurement to be 0.833 M ATP per $\mathrm{A}_{600\mathrm{nm}}$ per hour per protein fraction (see Table N3 of (1), the transporter LacY is approximately one-tenth of the total ∼2% protein fraction dedicated to fermentation). Assuming a cell volume of 2 fL (2), a dry cell mass of 0.278 pg (5), and 0.33 grams of dry cell mass per liter per unit of optical density (7), we convert this to be in terms of intracellular ATP flux: 351 M ATP per hour per protein fraction. We now must normalize this value to a concentration driving force in order to obtain $\hat{\nu}$. This can only be roughly estimated, as we are approximating many metabolic pathways as a single reaction. We take our reference driving force as the steady-state concentration of fructose-1,6-bisphosphate, a major glycolysis metabolite, measured as $c_{\mathrm{F16B}} = 1.52 \times 10^{-2}$ M in glucose-fed *E. coli* (see Table S3 of (8)). To convert from a flux of ATP to a flux of substrate, we divide by the $\Upsilon$ specified above. We also divide by the value of total amino acid content of the cell to yield $\hat{\nu} = 1 \times 10^{-6} \left(\frac{1}{\mathrm{h \cdot AA}}\right)$.

We now estimate $\hat{\beta}_i$, the constant relating intracellular and extracellular metabolite concentrations and the fraction of enzyme spent of transport enzymes to transport flux into of the cell. We assume all $\hat{\beta}_i$ are equal such that $\hat{\beta}_i \equiv \hat{\beta}$. As a reference enzyme, we use the LacY transporter of *E. coli*. We begin with the turnover number of LacY as measured by (9): 14 lactose per LacY per second at saturation. Using the number of amino acids in LacY ($N = 417$ AAs) and taking the cell volume as 2 fL (2), we obtain an intracellular transport flux of $1 \times 10^{-7}$ M per hour per AA. Note that we are only accounting for the enzymes importing the electron donor, and assuming the expression of generic transporters such as porins is constant among strains. We now normalize by the measured $K_m$ of the LacY enzyme (this approach makes $\hat{\beta}$ the first coefficient in the Taylor expansion of the enzyme's Michaelis-Menten kinetics). From (10) this is $K_m = 5 \times 10^{-4}$ M, giving us a final value of $\hat{\beta} = 2 \times 10^{-4} \left(\frac{1}{\mathrm{h \cdot AA}}\right)$.

Finally, we estimate the intracellular toxicity scale, $c^*$. This can only be approximately roughly, as there are many mechanisms by which intracellular metabolites can be detrimental to cell growth. As $c^*$ corresponds to the intracellular concentration at which growth rate is zero, we assume a value that is much higher than *E. coli*'s normal substrate levels. We choose this to be ten times the intracellular concentration of fructose-1,6-bisphosphate measured in (8), giving us $c^* = 1.52 \times 10^{-1}$ M.

Note that in this linear model of cell metabolism, the cell density reached within the chemostat is unconstrained and may reach physically unrealistic densities at high nutrient supply and low dilution rate. In a more realistic model, nonlinear effects would limit cell density. However, the linearized equations still allow comparison of growth rates of different strategies and thus are sufficient for our purposes of determining evolutionarily stable metabolic strategies.

**C. Non-dimensionalization.** In this section, we non-dimensionalize the dimensional equations into the equations presented in the main text. We first non-dimensionalize the enzyme budget by introducing $\hat{E}_i = E_i \xi B$ and $\hat{T}_i = T_i \xi B$. We then non-dimensionalize concentrations by the toxicity scale such that $\hat{c}_i = c_i c^*$. Finally, we non-dimensionalize time by normalizing to the timescale of intracellular metabolism $\xi B \hat{\nu}$ such that $\tau = t/(\xi B \hat{\nu})$. Substituting this into equations Eqs. S6-S8 yields:

$$\frac{dc_i}{dt} = -T_i \left(\frac{\hat{\beta}}{\hat{\nu}}\right)(c_i - c_{i,e}) + E_i(c_{i-1} - c_i/K_{\mathrm{eq},i}) - E_{i+1}(c_i - c_{i+1}/K_{\mathrm{eq},i+1}) - c_i g, \quad \text{[S9]}$$

$$\frac{dc_{i,e}}{dt} = \frac{\hat{s}_i}{c^* \xi B \hat{\nu}} + r_{\mathrm{v}} \rho T_i \left(\frac{\hat{\beta}}{\hat{\nu}}\right)(c_i - c_{i,e}) - \left(\frac{\hat{D}}{\xi B \hat{\nu}}\right) c_{i,e}, \quad \text{[S10]}$$

$$\frac{d\rho_\sigma}{dt} = -\rho_\sigma \left(\frac{\hat{D}}{\xi B \hat{\nu}}\right) + \rho_\sigma g, \quad \text{[S11]}$$

$$g = \left(1 - \sum_j c_j\right)\left(-\frac{\hat{g}_{\mathrm{m}}}{\xi B \hat{\nu}} + \sum_i \hat{\alpha}_i c^* \Upsilon_i E_i (c_{i-1} - c_i/K_{\mathrm{eq},i})\right). \quad \text{[S12]}$$

From this, we define new dimensionless parameters: $\beta = \hat{\beta}/\hat{\nu}$, $\alpha_i = \hat{\alpha}_i c^* \Upsilon_i$, $s_i = \frac{\hat{s}_i}{c^* \xi B \hat{\nu}}$, and $D = \frac{\hat{D}}{\xi B \hat{\nu}}$, and $g_{\mathrm{m}} = \frac{\hat{g}_{\mathrm{m}}}{\xi B \hat{\nu}}$. The final non-dimensionalized equations are thus:

$$1 = \sum_i E_i + \sum_i T_i, \quad [S13]$$
$$J_{c,i} = E_i(c_{i-1} - c_i/K_{\mathrm{eq},i}), \quad [S14]$$
$$J_{t,i} = T_i\beta(c_i - c_{i,e}), \quad [S15]$$
$$\Omega = 1 - \sum_i c_i, \quad [S16]$$
$$g = \Omega\left(-g_{\mathrm{m}} + \sum_i \alpha_i J_{c,i}\right), \quad [S17]$$
$$\frac{dc_i}{dt} = -J_{t,i} + J_{c,i} - J_{c,i+1} - gc_i, \quad [S18]$$
$$\frac{dc_{i,e}}{dt} = s_i + r_{\mathrm{V}}\rho J_{t,i} - Dc_{i,e}, \quad [S19]$$
$$\frac{d\rho_\sigma}{dt} = (g - D)\rho_\sigma. \quad [S20]$$

**Table S1. List of dimensional and dimensionless parameters, see "Model parameterization" section for details.**

| *Parameter* | *Meaning* | *Value* | *Units* |
|---|---|---|---|
| $\xi$ | Fraction of proteome budget available for transport and metabolism | 0.1 | unitless |
| $B$ | Proteome budget | $1.9 \times 10^9$ | amino acids per cell |
| $\hat{\nu_i}$ | Rate coefficient of metabolic enzymes | $1 \times 10^{-6}$ | $\frac{1}{\mathrm{h}\cdot\mathrm{AA}}$ |
| $\hat{\beta}_i$ | Rate coefficient of transport enzymes | $2 \times 10^{-4}$ | $\frac{1}{\mathrm{h}\cdot\mathrm{AA}}$ |
| $K_{\mathrm{eq},i}$ | Metabolic reaction equilibrium constant | varies | |
| $c^*$ | Concentration scale of growth penalty | $1.5 \times 10^{-1}$ | M |
| $\hat{\alpha}_i$ | Conversion factor between ATP flux and growth rate | 0.1 | $\mathrm{M}^{-1}$ |
| $\Upsilon$ | ATP yield per molecule substrate | 12 | unitless |
| $\hat{s}_i$ | Substrate inflow rate | varies | M/h |
| $r_{\mathrm{V}}$ | Ratio of individual cell to chemostat volume | $2 \times 10^{-15}$ | unitless |
| $\hat{D}$ | Chemostat dilution rate | varies | |
| $\hat{g}_{\mathrm{m}}$ | Maintenance growth rate requirement | $10^{-1}$ | $\mathrm{h}^{-1}$ |
| $\beta$ | Dimensionless transporter rate coefficient | $2 \times 10^2$ | unitless |
| $\alpha$ | Dimensionless conversion factor between substrate flux and growth rate | $1.8 \times 10^{-1}$ | unitless |
| $s_i$ | Dimensionless substrate inflow rate | varies | unitless |
| $D$ | Dimensionless dilution rate | varies | unitless |
| $g_{\mathrm{m}}$ | Dimensionless maintenance growth rate requirement | $5 \times 10^{-4}$ | unitless |

### 2. Numerical methods

**A. Overview of simulation methods.** All numerical integration is performed using MATLAB's *ode15s* function. Steady states are identified using the relative ratio of the derivative at a given point to the current value of the state variable, i.e. $y'(t)/y(t) < \epsilon_{\mathrm{s}}$, where $\epsilon_{\mathrm{s}} = 10^{-9}$.

To compute evolutionarily stable consortia in a given condition, we begin by initializing the abiotic environment and allow it to come to steady state. We then identify the strain with the highest invasion growth rate. If this growth rate is above the dilution rate, we insert the strain into the simulation and allow the new ecosystem to reach steady state. We then repeat the invasion procedure until no new strains can invade. Below, we further detail the methods used to identify and invade new strains into the ecosystem.

**B. Strain invasion methods.** To identify the next invading strain, we perform a numerical optimization procedure to find the strain with the highest invasion growth rate within the current environment. To compute this growth rate for a given strain, we assume an infinitesimal population size of the invader and compute the steady state of the strain's internal metabolism given fixed external conditions set by the current environment in the chemostat (i.e. $c_{i,e} = \mathrm{const}$). We consider only the P1, P2, and C strain classes. For each strain class, we derived analytical expressions for the steady state in the absence of growth (i.e., neglecting internal dilution due to growth) and use these in our optimization procedure. We compared this approach for computing growth rates to one in which we compute the invading strain's internal metabolism via numerical integration without neglecting growth dilution and found they lead to similar final consortia. In the few cases where the final consortia produced by these two approaches differ, this occurs due to the emergence of strains relying entirely on growth dilution to eliminate waste products. We consider such strains to be pathological and not reflective of realistic metabolic strategies.

Optimization is performed via the *fmincon* constrained optimization function within MATLAB, with the solution constrained to obey the enzyme budget constraint. To ensure that we identify the best possible invading strain, we perform the optimization many times for a given invasion: (1) across different initial enzyme strategy values, (2) using different optimization constraints, e.g. some runs restricted only to finding P1 strains, and (3) using either the 'interior-point' or 'sqp' algorithm. From all of these optimizations, the strategy with the highest invasion growth rate is selected for possible invasion. Given the complexity

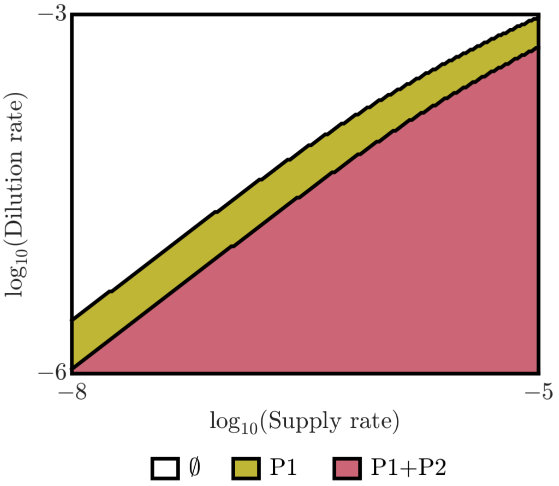

**Fig. S1.** Phase diagram of evolutionarily stable communities in the thermodynamic toxicity model with no energy yield from the second metabolic step ($\alpha_2 = 0$) and $K_{\mathrm{eq},i} = 1$, as a function of nutrient supply rate, $s_0$, and dilution rate, $D$. Model parameters can be found in *SI Appendix* 1 and aside from $\alpha_2$ are equivalent to Fig. 2A. "∅" indicates environments where no strains can persist.

and high-dimensionality of the invasion growth function, we cannot guarantee that this procedure always finds the optimal strain. However, we find that in practice many of these above optimization runs produce nearly-identical optimal strains and that the algorithm's outputs are in agreement with available analytical results.

We add the new strain to the simulation if its initial invasion growth rate exceeds the dilution rate by a given tolerance $g_{\mathrm{inv}} > D(1+\epsilon_{\mathrm{g}})$, where $\epsilon_{\mathrm{g}} = 10^{-3}$ in our simulations. This finite tolerance is required to prevent invasion of many nearly-identical strains. If the invading strain is from a new class (e.g. P2 in a system that currently contains no P2), it is added to the simulation with an invasion biomass of $10^{11}$ cells/L. If the strain is from an existing class (e.g. a different strain of P1) we remove the existing strain of that class and replace it with the new strain with an abundance equivalent to that of the old strain. We find that this replacement procedure does not impact the resulting evolutionarily stable state, but does prevent pathological cases of many nearly-identical strains of the same class coexisting in a nearly-stable pseudo-equilibrium.

### 3. Thermodynamic toxicity model with zero energy yield from second metabolic reaction

In this section, we show that similar evolutionarily stable strategies emerge in a model in which the second metabolic step yields no energy (i.e. $\alpha_2 = 0$). In Fig. S1, we show a numerically generated phase diagram equivalent to Fig. 2A except with $\alpha_2 = 0$. As can be seen, the P1 + P2 consortium still emerges despite there being no energetic benefit to further metabolizing $c_1$ into $c_2$. This occurs because while performing the second metabolic step does not produce energy, it allows P2 to avoid being adversely affected by the extracellular concentration of $c_1$, thus enabling P2 to invade a P1 community that has polluted its environment with $c_1$.

### 4. Thermodynamic toxicity model with varying $K_{\mathrm{eq},i}$

In this section, we show that similar evolutionarily stable strategies emerge in a model in which the $K_{\mathrm{eq},i}$ differ between the two reactions. In Fig. S2, we show a numerically generated phase diagram equivalent to Fig. 2A except with $K_{\mathrm{eq},1} = 1$ and $K_{\mathrm{eq},2} = 0.1$. As can be seen, the P1 + P2 consortium still emerges despite the second reaction being less thermodynamically favorable. As in the case of zero energy yield from the second reaction, this occurs because performing the second metabolic step allows P2 to avoid being adversely affected by the extracellular concentration of $c_1$. This enables P2 to invade a P1 community that has polluted its environment with $c_1$ despite the lower thermodynamic favorability of the second reaction.

### 5. Derivation of evolutionarily stable strategies in a simplified thermodynamic model

In this section, we derive the evolutionarily stable strategies in a slightly simplified thermodynamic model. In this version, we neglect the role of intracellular metabolite dilution due to growth (i.e. the term $gc_i$ in Eq. S18). We find this simplification does not substantially alter the evolutionarily stable states nor the phase boundaries (see Fig. S3), but does alter the relative abundance of evolutionarily stable strategies in the final community composition.

We calculate the evolutionarily stable strategy or strategies for each type of consortium (e.g. P1 alone or P1 + P2). To do so, we first analytically derive the external steady-state nutrient concentration of the input nutrient as a function of the enzyme strategies and the other model parameters $\hat{\theta}$, including supply rate and dilution:

$$c_{0,e}^{*} = f(E_i, T_i, \hat{\theta}). \qquad [\mathrm{S21}]$$

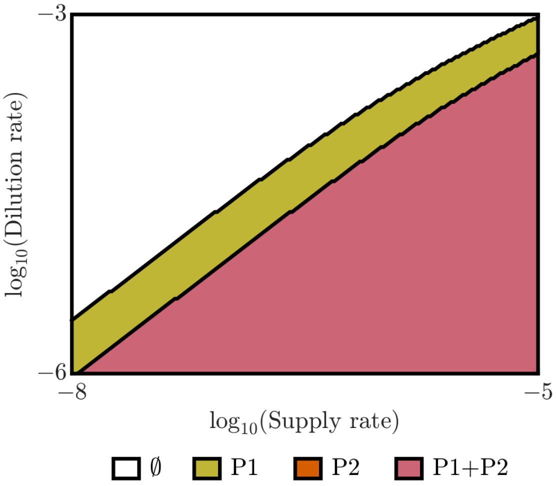

**Fig. S2.** Phase diagram of evolutionarily stable communities in the thermodynamic toxicity model with each energy-yielding reaction having different $K_{\text{eq},i}$ ($K_{\text{eq},1} = 1$, $K_{\text{eq},2} = 0.1$). The diagram is shown as a function of nutrient supply rate, $s_0$, and dilution rate, $D$. Model parameters can be found in *SI Appendix* 1 and aside from $K_{\text{eq},2}$ are equivalent to Fig. 2A. "∅" indicates environments where no strains can persist.

The competitive ability of a consortium is determined by its ability to deplete this nutrient and create the harshest possible environment. We thus use the method of Lagrange multipliers to find the set of enzyme strategies $E_i$ and $T_i$ that minimize $c^*_{0,e}$. The constraint equation to ensure the enzyme strategies sum to the budget is

$$h(E_i, T_i) = -1 + \sum_i E_i + \sum_i T_i = 0. \quad \text{[S22]}$$

Thus, the Lagrangian function is

$$\mathcal{L}(E_i, T_i, \hat{\theta}) = c^*_{0,e}(E_i, T_i, \hat{\theta}) - \lambda h(E_i, T_i), \quad \text{[S23]}$$

where $\lambda$ is the Lagrange multiplier and our optimization problem is to find

$$\nabla_{E_i,T_i,\lambda}\mathcal{L}(E_i, T_i, \hat{\theta}) = 0. \quad \text{[S24]}$$

The $E_i$ and $T_i$ satisfying these equations are the evolutionarily stable strategies. Below, we show this calculation in full only for the evolutionarily stable P1 strategy and only report final results for the other consortia, as the calculations are mechanically very similar. In practice, we implement the bulk of these calculations using MATLAB's symbolic algebra toolbox and these scripts are available within the manuscript's code repository.

**A. Evolutionarily stable strategies of P1 alone.** A community of P1 growing alone is governed by the following dynamical equations:

$$1 = E_1 + T_0 + T_1, \quad \text{[S25]}$$

$$J_{c,1} = E_0(c_0 - c_1/K_{\text{eq},1}), \quad \text{[S26]}$$

$$J_{t,0} = T_0\beta(c_0 - c_{0,e}), \quad \text{[S27]}$$

$$J_{t,1} = T_1\beta(c_1 - c_{1,e}), \quad \text{[S28]}$$

$$g = \alpha_1 J_{c,1} - g_{\text{m}}, \quad \text{[S29]}$$

$$\frac{dc_0}{dt} = -J_{t,0} - J_{c,1}, \quad \text{[S30]}$$

$$\frac{dc_1}{dt} = -J_{t,1} + J_{c,1}, \quad \text{[S31]}$$

$$\frac{dc_{0,e}}{dt} = s_0 + r_{\text{v}}\rho J_{t,0} - Dc_{0,e}, \quad \text{[S32]}$$

$$\frac{dc_{1,e}}{dt} = r_{\text{v}}\rho J_{t,1} - Dc_{1,e}, \quad \text{[S33]}$$

$$\frac{d\rho}{dt} = (g - D)\rho. \quad \text{[S34]}$$

Given these equations, the steady-state concentration of compound 0 is

$$c_{0,e}^* = \frac{D(D+g_\mathrm{m})E_1T_0 + D(D+g_\mathrm{m})E_1K_{\mathrm{eq},1}T_1 + D(D+g_\mathrm{m})K_{\mathrm{eq},1}T_0T_1\beta + E_1s_0T_0T_1\alpha_1\beta}{DE_1T_0T_1\alpha_1\beta(K_{\mathrm{eq},1}+1)}, \quad \text{[S35]}$$

and thus the Lagrangian is $\mathcal{L}(E_i, T_i, \hat{\theta}) = c_{0,e}^*(E_i, T_i, \hat{\theta}) - \lambda(-1 + E_1 + T_0 + T_1)$. Taking partial derivatives of the Lagrangian with respect to the enzyme levels yields

$$\frac{\partial \mathcal{L}}{\partial E_1} = \frac{-(D+g_\mathrm{m})K_{\mathrm{eq},1}}{E_1^2\alpha_1(K_{\mathrm{eq},1}+1)} - \lambda, \quad \text{[S36]}$$

$$\frac{\partial \mathcal{L}}{\partial T_0} = \frac{-(D+g_\mathrm{m})K_{\mathrm{eq},1}}{T_0^2\alpha_1\beta(K_{\mathrm{eq},1}+1)} - \lambda, \quad \text{[S37]}$$

$$\frac{\partial \mathcal{L}}{\partial T_1} = \frac{-(D+g_\mathrm{m})}{T_1^2\alpha_1\beta(K_{\mathrm{eq},1}+1)} - \lambda. \quad \text{[S38]}$$

$$\frac{\partial \mathcal{L}}{\partial \lambda} = -1 + E_1 + T_0 + T_1. \quad \text{[S39]}$$

We set these equations to zero to find the optimum. Equating the right-hand sides of Eqs. S36-S38 to one another and simplifying provides the relative relationships between the enzyme levels (e.g., $T_0 = E_1/\sqrt{\beta}$) and these relationships can then be substituted into Eq. S39 to compute the evolutionarily stable strategies:

$$E_1 = \frac{1}{1 + \frac{1}{\sqrt{\beta}} + \frac{1}{\sqrt{\beta K_{\mathrm{eq},1}}}}, \quad \text{[S40]}$$

$$T_0 = \frac{1}{1 + \sqrt{\beta} + \frac{1}{\sqrt{K_{\mathrm{eq},1}}}}, \quad \text{[S41]}$$

$$T_1 = \frac{1}{1 + \sqrt{K_{\mathrm{eq},1}} + \sqrt{\beta K_{\mathrm{eq},1}}}. \quad \text{[S42]}$$

To determine the steady-state value of $c_{0,e}^*$, one can then substitute the above enzyme levels into S35.

**B. Evolutionarily stable strategy of P2 alone.** For P2 alone, repeating the above Lagrangian calculation procedure leads to:

$$E_1 = \frac{1}{1 + \frac{1}{\sqrt{\beta}} + \frac{1}{\sqrt{K_{\mathrm{eq},1}}} + \frac{1}{\sqrt{K_{\mathrm{eq},1}K_{\mathrm{eq},2}\beta}}}, \quad \text{[S43]}$$

$$T_0 = \frac{1}{1 + \sqrt{\beta} + \frac{1}{\sqrt{K_{\mathrm{eq},1}K_{\mathrm{eq},2}}} + \sqrt{\frac{\beta}{K_{\mathrm{eq},1}}}}, \quad \text{[S44]}$$

$$E_2 = \frac{1}{1 + \sqrt{K_{\mathrm{eq},1}} + \frac{1}{\sqrt{K_{\mathrm{eq},2}\beta}} + \sqrt{\frac{K_{\mathrm{eq},1}}{\beta}}}, \quad \text{[S45]}$$

$$T_2 = \frac{1}{1 + \sqrt{K_{\mathrm{eq},1}K_{\mathrm{eq},2}} + \sqrt{K_{\mathrm{eq},2}\beta} + \sqrt{K_{\mathrm{eq},1}K_{\mathrm{eq},2}\beta}}. \quad \text{[S46]}$$

**C. Evolutionarily stable strategy of P1 + P2 and P1 + C consortia.** For consortia, the optimization calculation is performed with two Lagrange multipliers, one corresponding to each strain's enzyme budget constraints. For example, the form of the Lagrangian for P1 + C will be

$$\mathcal{L} = c_{0,e}^*(E_1^{\mathrm{P1}}, T_0^{\mathrm{P1}}, T_1^{\mathrm{P1}}, E_2^{\mathrm{C}}, T_1^{\mathrm{C}}, T_2^{\mathrm{C}}, \hat{\theta}) - \lambda^{\mathrm{P1}}(-1 + E_1^{\mathrm{P1}} + T_0^{\mathrm{P1}} + T_1^{\mathrm{P1}}) - \lambda^{\mathrm{C}}(-1 + E_2^{\mathrm{C}} + T_1^{\mathrm{C}} + T_2^{\mathrm{C}}). \quad \text{[S47]}$$

Despite this difference, the calculation proceeds very similarly, as each constraint term only depends on the enzyme levels of a single strain. Solving for the zero point of the Lagrangian derivatives decouples into two separate systems of equations, one for each strain. In both the P1 + P2 and P1 + C consortia, the stable P1 and P2 members of the consortia are identical to stable strategies in the P1 alone and P2 alone cases shown above. The stable C strategy in the P1 + C consortium is similar in form to the stable P1 strategy:

$$E_1^{\mathrm{C}} = \frac{1}{1 + \frac{1}{\sqrt{\beta}} + \frac{1}{\sqrt{\beta K_{\mathrm{eq},2}}}}, \quad \text{[S48]}$$

$$T_1^{\mathrm{C}} = \frac{1}{1 + \sqrt{\beta} + \frac{1}{\sqrt{K_{\mathrm{eq},2}}}}, \quad \text{[S49]}$$

$$T_2^{\mathrm{C}} = \frac{1}{1 + \sqrt{K_{\mathrm{eq},2}} + \sqrt{\beta K_{\mathrm{eq},2}}}. \quad \text{[S50]}$$

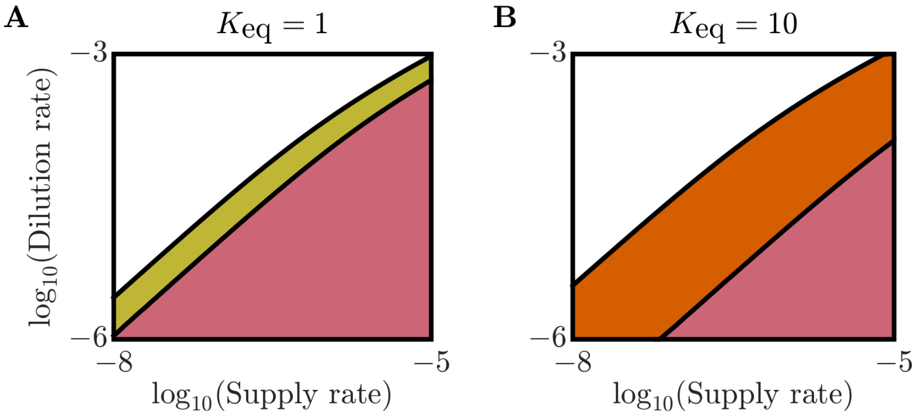

**Fig. S3.** Version of Fig. 2AB generated from the simplified thermodynamic toxicity model in which the intracellular dilution term $gc_i$ is neglected. All model parameters the same as in Fig. 2AB. The evolutionarily stable consortia is determined by minimizing the expression for the steady-state $c_{0,e}$ of each consortia type and identifying the consortia that leads to the lowest $c_{0,e}$ in a given condition. As can be seen, the phase diagram of this simplified model is similar to that of the full model, validating our analytical approximation.

**D. Determination of cross-feeding stability.** To determine whether cross-feeding is ever evolutionarily stable in the simplified thermodynamic model, we substitute in the optimal strategies of P1 + P2 and P1 + C into their respective expressions for $c^*_{0,e}$. We then compute the difference between these two minimum $c^*_{0,e}$ values:

$$c_{0,e}^{*,\mathrm{P1+C}} - c_{0,e}^{*,\mathrm{P1+P2}} = \frac{(D+g_{\mathrm{m}})\left(1+\sqrt{K_{\mathrm{eq},2}}+\sqrt{\beta K_{\mathrm{eq},2}}\right)^2}{\alpha_2\beta(K_{\mathrm{eq},1}+K_{\mathrm{eq},1}K_{\mathrm{eq},2}+1)} + \frac{(D+g_{\mathrm{m}})K_{\mathrm{eq},2}\left(1+\sqrt{K_{\mathrm{eq},1}}+\sqrt{\beta K_{\mathrm{eq},1}}\right)^2}{\alpha_1\beta(K_{\mathrm{eq},1}+K_{\mathrm{eq},1}K_{\mathrm{eq},2}+1)} - \frac{(D+g_{\mathrm{m}})\left(1+\sqrt{\beta K_{\mathrm{eq},2}}+\sqrt{K_{\mathrm{eq},1}K_{\mathrm{eq},2}}+\sqrt{\beta K_{\mathrm{eq},1}K_{\mathrm{eq},2}}\right)^2}{(\alpha_1+\alpha_2)\beta(K_{\mathrm{eq},1}+K_{\mathrm{eq},1}K_{\mathrm{eq},2}+1)}. \quad [S51]$$

In order for cross-feeding to be stable, there must be a regime where $c_{0,e}^{*,\mathrm{P1+C}} - c_{0,e}^{*,\mathrm{P1+P2}} < 0$, as this would correspond to the scenario where the optimal P1 + C consortium can create an environment that has such low nutrient levels that the optimal P1 + P2 consortium cannot invade. For the case of equal $K_{\mathrm{eq},i}$ and equal $\alpha_i$, we use the *isAlways* function in MATLAB's symbolic algebra toolbox to analytically prove that this expression is never negative and thus cross-feeding is never evolutionarily stable in this case.

For the case of unequal $K_{\mathrm{eq},i}$ and unequal $\alpha_i$, we were unable to prove or disprove whether the difference is always positive via symbolic algebra or manual calculation approaches. These approaches were similarly inconclusive for the case of unequal $K_{\mathrm{eq},i}$ with equal $\alpha_i$ and vice versa. To probe this general case, we turned to numerical evaluation of Eq. S51. As this is a single algebraic expression, we can test a much larger range of parameters than is possible with our full evolutionary simulation. As $D + g_{\mathrm{m}}$ simply rescales the magnitude of the difference, we set $D + g_{\mathrm{m}} = 1$ without loss of generality. For $\alpha_1$, $\alpha_2$, $K_{\mathrm{eq},1}$, $K_{\mathrm{eq},2}$, and $\beta$, we consider a range of 100 logarithmically spaced values between $10^{-5}$ and $10^5$. We consider all combinations of these parameters for a total of $1 \times 10^{10}$ combinations. Across all of these tests, we do not find a negative value of $c_{0,e}^{*,\mathrm{P1+C}} - c_{0,e}^{*,\mathrm{P1+P2}}$.

### 6. Bistability in the osmotic toxicity model

We find that the osmotic toxicity model can exhibit bistability when environmental toxicity is strong. To demonstrate this behavior, we show an example in Fig. S5 of a P1+P2 consortium that can only successfully establish at sufficiently high population density. When this particular P2 strain is introduced on its own at a density of $7 \times 10^{11}$ cells/L, the extracellular concentration of $c_{0,e}$, which is toxic to the cell's metabolism, increases from its initial value and drives the P2 strain to extinction. The same occurs when a small amount of the P1 strain is added into the system. However, once a sufficiently large amount of P1 is introduced, the P1+P2 consortium is able to collectively detoxify the environment by stopping the increase in $c_{0,e}$, thus allowing the consortium to stably colonize the environment.

In the evolutionary algorithm, we introduce new strains at a sufficiently high abundance ($10^{11}$ cells/L for new strain classes) such that we have not observed any instances of this bistable extinction behavior influencing the outcome of our evolution simulations. To determine whether other forms of bistability might influence our computed evolutionary stable states, we performed tests to detect possible hysteresis near evolutionary transitions. In these numerical experiments, we first determine the evolutionarily stable community in an environmental condition near a phase boundary. We then alter the environment and re-run the evolution procedure, now starting from the evolutionarily stable consortia of the old environment rather than starting from an abiotic environment. We perform these changes in environmental parameter stepwise across a phase boundary, and then perform the same steps in reverse to cross the phase boundary from the opposite direction. None of these tests detected hysteresis, and the states reached by the stepwise procedure were the same as those reached when starting from an

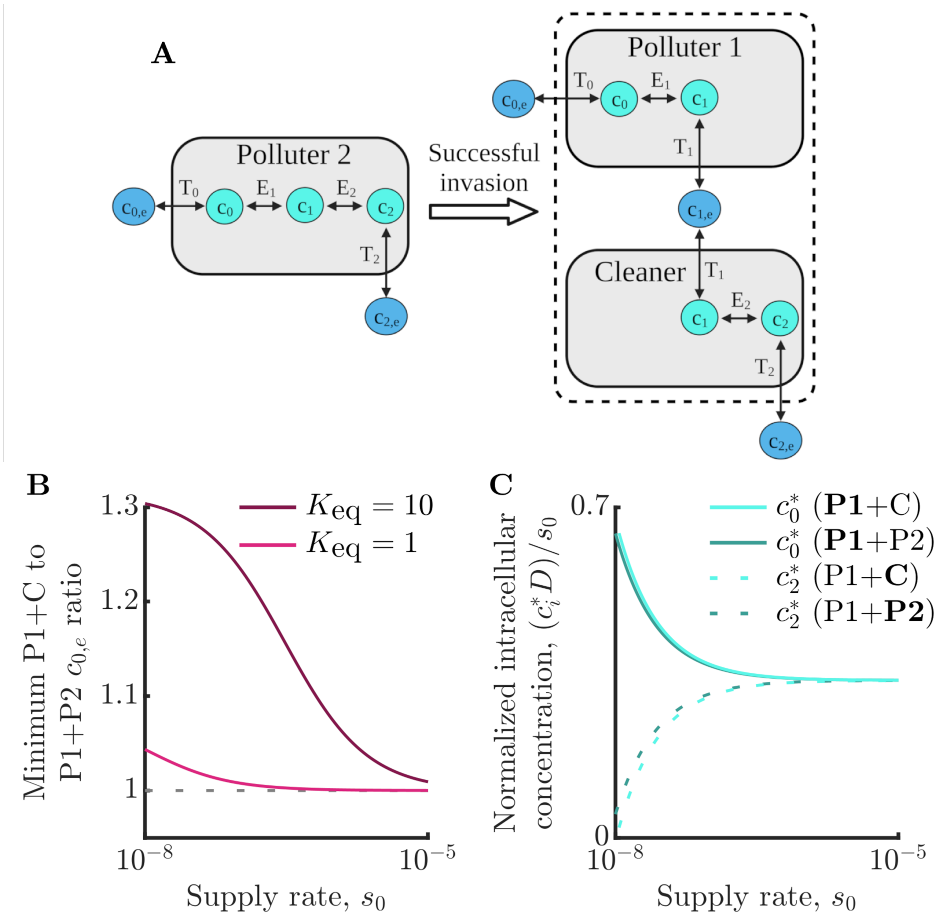

**Fig. S4.** Version of Fig. 3 generated analytically from simplified thermodynamic toxicity model in which the intracellular dilution term $gc_i$ is neglected. All model parameters the same as in Fig. 3. As can be seen, the $c_{0,e}$ ratio in the simplified model is similar to that of the full model, validating our analytical approximation.

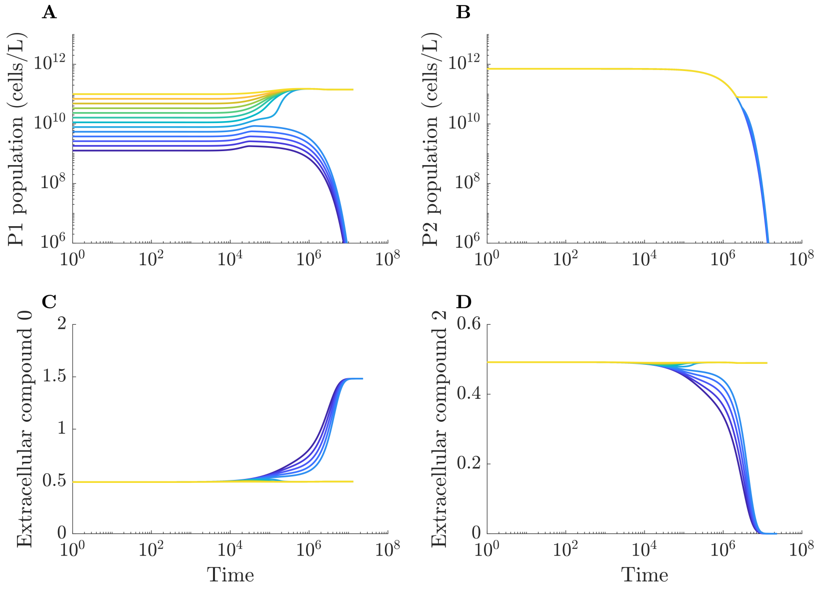

**Fig. S5.** Example of bistability in the osmotic toxicity model. Different colored trajectories represent simulations with different starting abundances of P1 and the same initial abundance of P2. These simulations lead to one of two distinct outcomes: either coexistence of P1 and P2 or total extinction. Simulation run with $D = 10^{-6}$, $s_0 = 1.5 \times 10^{-6}$, $[E_1^{P1}, E_2^{P1}, T_0^{P1}, T_1^{P1}, T_2^{P1}] = [0.87, 0, 0.06, 0.07, 0]$, $[E_1^{P2}, E_2^{P2}, T_0^{P2}, T_1^{P2}, T_2^{P2}] = [0, 0.88, 0, 0.04, 0.08]$, $\rho_{P2}(0) = 7 \times 10^{11}$ cells/L, $[c_{1,e}(0), c_{2,e}(0), c_{3,e}(0)] = [0.49, 0.49, 0]$, $[c_0^{P1}(0), c_1^{P1}(0), c_2^{P1}(0)] = [0, 0.49, 0.49]$, and $[c_0^{P2}(0), c_1^{P2}(0), c_2^{P2}(0)] = [0.49, 0.49, 0.49]$. All other parameters same as in main text simulations.

abiotic environment. We show an example in Fig. S8. As can be seen, the evolutionarily stable states resulting from stepwise alteration of the external environment ('Stepwise') are the same as those computed from an abiotic initial condition ('Abiotic').

Our algorithm for identifying evolutionarily stable consortia is not meant to model actual evolutionary trajectories, but how might this bistability phenomenon influence more realistic evolutionary trajectories? The examples of bistability we observe only occur when initial conditions are far from ecological steady states. Indeed, we initially identified the example we show in Fig. S5 when the resident P1 of a stable P1+P2 consortium was entirely removed and replaced with a slightly different P1 strain at much lower abundance. Thus, this bistability may influence the initial colonization of an environment by a community, representing an Allee effect in which a minimum population is required for colonization. We hypothesize that once a stable community is established, mutations or invasions will be unlikely to lead to mass extinction. More detailed study of the role of such bistability in cross-feeding evolution represents an interesting area of future study.

### 7. Thermodynamic toxicity model with Monod growth kinetics

The models we analyze in the main text assume that the catalysis flux $J_{c,i}$ is a purely linear function of intracellular concentrations. In this section, we show that the addition of more realistic saturating growth kinetics still does not allow for the evolution of cross-feeding in the thermodynamic toxicity model and results in similar evolutionarily stable strategies. In

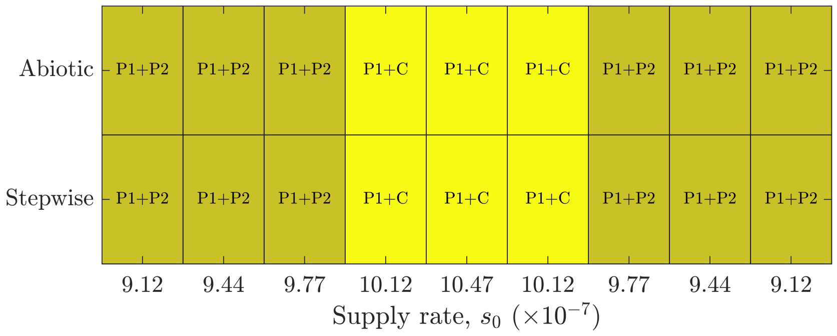

**Fig. S6.** Example of a test for hysteresis near a phase boundary of the osmotic toxicity model. Simulations performed by varying supply rate $s_0$ at constant dilution rate $D = 10^{-6}$, with all other parameters the same as in Fig. 4B. The 'Abiotic' row represents the evolutionarily stable states reached when the evolutionarily algorithm is initialized from an abiotic state at the specified environmental conditions. The 'Stepwise' row represents the state reached from stepwise environmental changes, with the evolutionarily algorithm being initialized using the evolutionarily stable consortium from the previous environment.

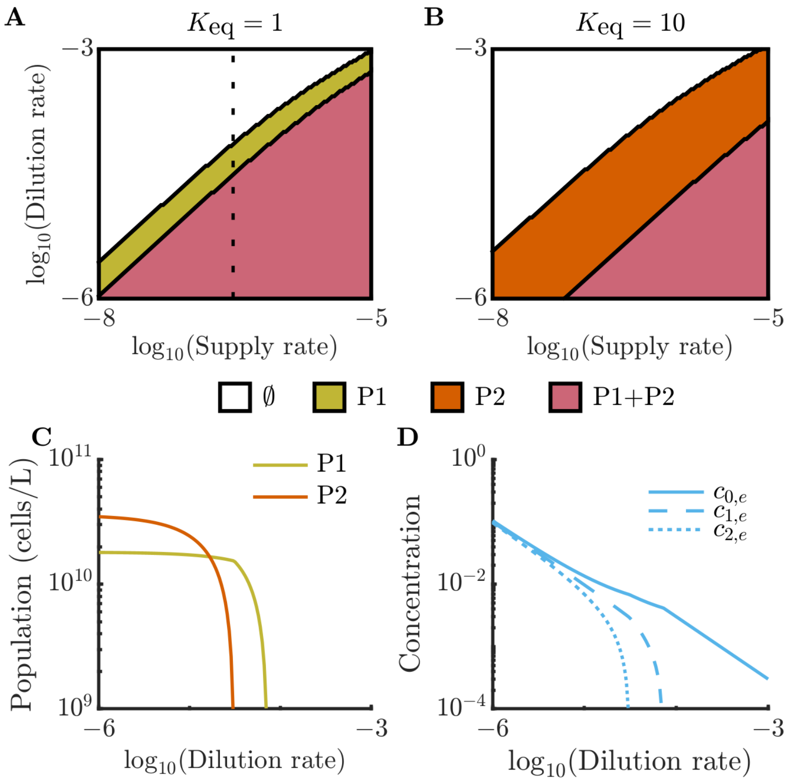

**Fig. S7.** Version of Fig. 2 with a saturating growth function. Model parameters can be found in *SI Appendix* 1 and aside from the saturating growth function (Eq. S52) with Monod constants $K_{i-1} = K_i = 1$, parameters are the same as in Fig. 2.

particular, the growth function we consider is a generalization of the classical Michaelis-Menten function:

$$J_{c,i} = \frac{E_i(c_{i-1} - c_i/K_{\rm eq})}{1 + c_{i-1}/K_{i-1} + c_i/K_i}, \qquad \text{[S52]}$$

where the $K_i$ are saturation constants (11). We show an example of the resulting evolutionarily stable strategies in Fig. S7. As can be seen, this phase diagram is qualitatively similar to Fig. 2A, which is computed with the same parameters with the exception of the growth function. We extensively numerically explored the version of the model with growth saturation and did not identify any parameter regions with evolutionarily stable cross-feeding. Our intuition for this result is that while the saturating growth function is nonlinear, it is still monotonic in the primary substrate, and as a result there is no regime in which intracellular toxicity makes it favorable to split metabolism between two strains.

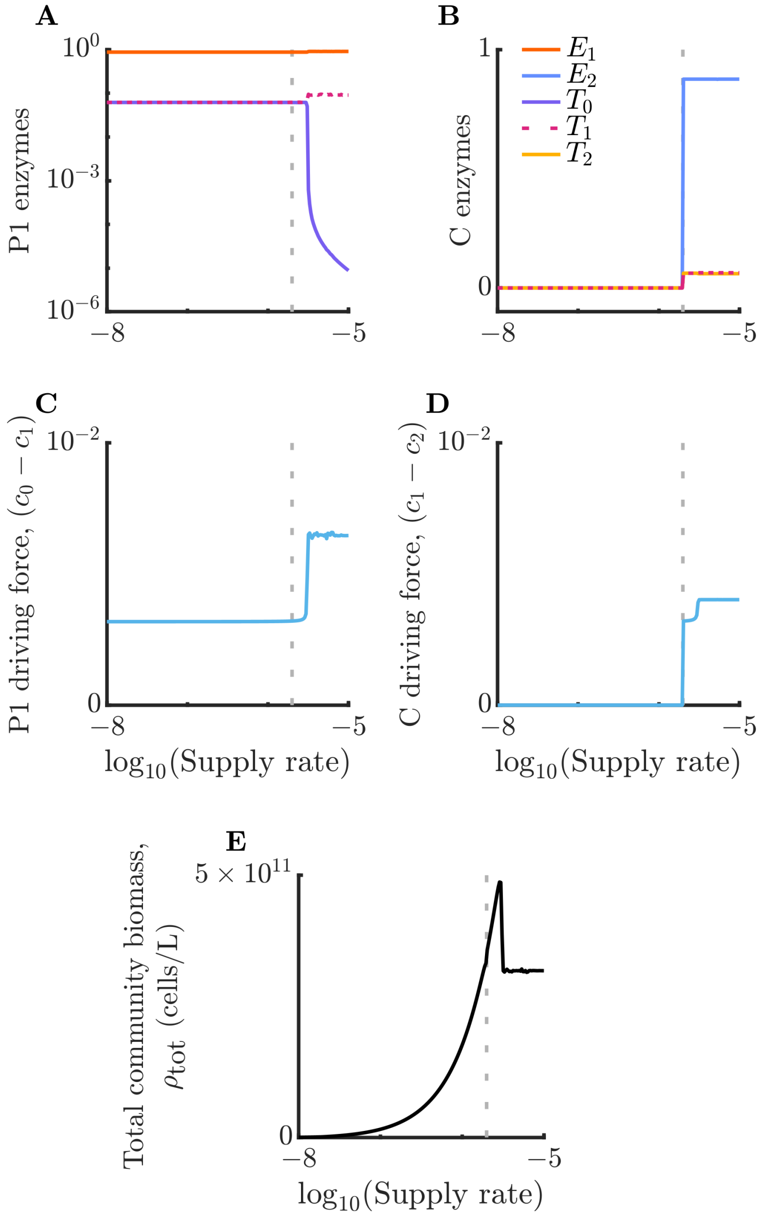

**Fig. S8.** Community properties as a function of supply rate, corresponding to the communities shown in Fig. 4DE. The vertical dashed line corresponds to the onset of the cross-feeding phase. These plots shows the enzyme strategy of P1 (*A*), enzyme strategy of C (*B*), metabolic driving force of P1 (*C*), metabolic driving force of C (*D*), and total community biomass (*E*). As can be seen, these quantities change abruptly, albeit continuously, after the initial entry into the cross-feeding phase, corresponding to the onset of the pessimization regime.
